# Data structures based on *k*-mers for querying large collections of sequencing datasets

**DOI:** 10.1101/866756

**Authors:** Camille Marchet, Christina Boucher, Simon J Puglisi, Paul Medvedev, Mikaël Salson, Rayan Chikhi

## Abstract

High-throughput sequencing datasets are usually deposited in public repositories, e.g. the European Nucleotide Archive, to ensure reproducibility. As the amount of data has reached petabyte scale, repositories do not allow to perform online sequence searches; yet such a feature would be highly useful to investigators. Towards this goal, in the last few years several computational approaches have been introduced to index and query large collections of datasets. Here we propose an accessible survey of these approaches, which are generally based on representing datasets as sets of *k*-mers. We review their properties, introduce a classification, and present their general intuition. We summarize their performance and highlight their current strengths and limitations.

## INTRODUCTION

Over the past decade the cost of sequencing has decreased dramatically, making the generation of sequence data more accessible. This has led to increasingly ambitious sequencing projects. For example, the 1,000 Genomes Project, which began in 2008 and completed in 2012 (Clarke et al., 2012), led to the 100,000 Genomes Project, which began in 2014 and completed in 2018 (Turnbull et al., 2018). There are dozens of other large-scale sequencing projects completed or underway, including GEUVADIS (Lappalainen et al., 2013), GenomeTrakr (Timme et al., 2018), and MetaSub (The MetaSUB International Consortium, 2016). An overwhelming amount of public data is now available at EBI’s European Nucleotide Archive (ENA) (Cook et al., 2018) and NCBI’s Sequence Read Archive (SRA) (Leinonen et al., 2010). The possibility of analyzing these collections of datasets, alone or in combination, creates vast opportunities for scientific discovery, exceeding the capabilities of traditional laboratory experiments. For this reason, there has been a substantial amount of work in developing methods to store and compress collections of high-throughput sequencing datasets in a manner that supports various queries.

In this paper, we use the term <u>dataset</u> to refer to a set of reads resulting from sequencing an individual sample (e.g., DNA-seq, or RNA-seq, or metagenome sequencing). Sequencing is routinely performed not only on a single sample but on a collection of samples, resulting in a collection of datasets. For instance, 100,000 human genomes were sequenced for the 100,000 Genome Project and over 300,000 bacterial strains were sequenced for GenomeTrakr. One basic query that is fundamental to many different types of analyses of such collections of datasets can be formulated as follows: given a sequence, identify all datasets in which this sequence is found. For example, consider the problem of finding a RNA transcript within a collection of RNA-seq datasets. Similarly, we can ask to find which datasets contain a specific DNA sequence, such as a gene or a non-coding element, in a collection of bacterial strain genomes. In this paper, we present an overview of recent bioinformatics methods (Figure 1) created to handle these types of queries.

**Fig. 1.**
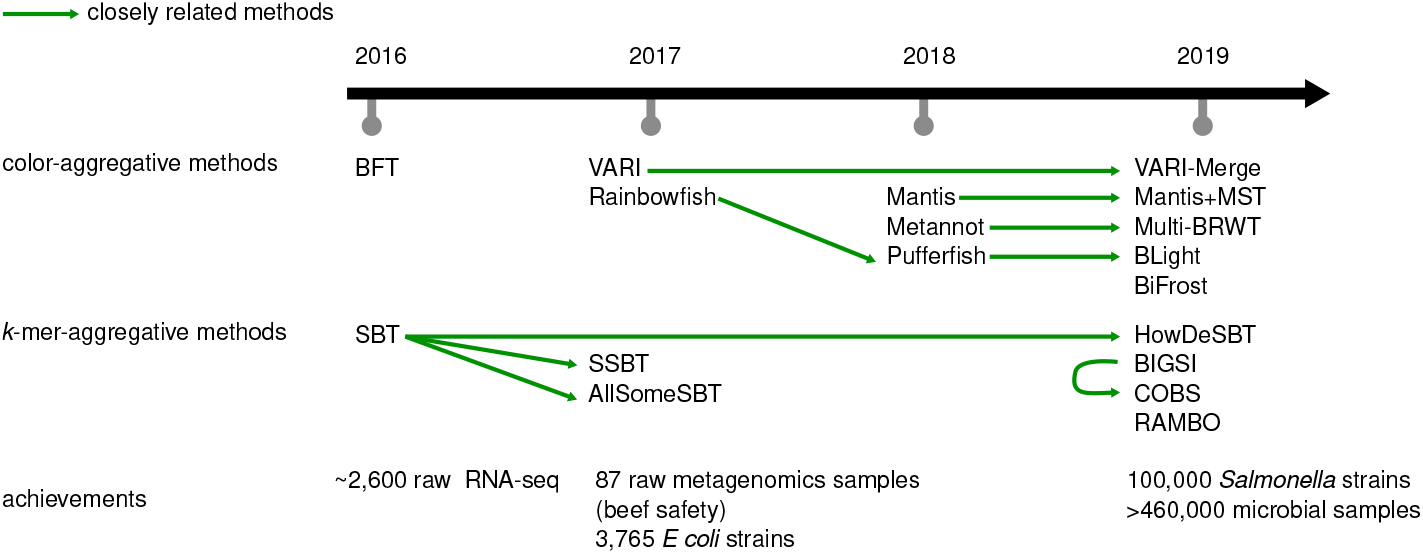
Timeline, main relationships and application highlights for the methods covered in this survey.

Given the size of many collections and datasets, several different paradigms for storing them so that they can be efficiently queried have been developed – many of which continue to be extended and explored. One paradigm is to store and index datasets as sets of *k*-length substrings, which are referred to as <u>*k*-mers</u>. We will refer to collections of datasets as <u>sets of *k*-mer sets</u>. The methods that use this paradigm build an index of all *k*-mer sets, and support the basic query described above by splitting the query sequence into *k*-mers and determining their presence or absence in the index.

As we will discuss in this survey, this paradigm has proven to be useful in several ways. First, sets of *k*-mers are a more concise representation of the set of sequences of the samples, as they abstract some of the redundancy inherent in high sequencing coverage. Second, genetic variation and sequencing errors can be dealt with in a more efficient, albeit less accurate way than using sequence alignment. Instead of performing inexact pattern matching, as aligners do, *k*-mer based methods can simply examine the fraction of matching *k*-mers within the query sequence. Since their initial development, *k*-mer based analysis approaches have been widely-adopted in the bioinformatics community due to their efficiency and ability to accurately summarize and compare large datasets. Hence, the methods we describe in this survey were imperative to a large number of biological analyses; from elucidating the evolutionary dynamics between substrains of *Staphylococcus aureus* (Young et al., 2012) to identifying of recombination hotspots in bird species (Singhal et al., 2015) or enabling the search of over 400,000 viral and bacterial species (Bradley et al., 2019b).

Even though *k*-mer based methods have been popular, there do exist some trade-offs. More specifically, storing data using a *k*-mer index comes with some loss of information since it only gives information for each constituent *k*-mer of a sequence – rather than about the entire sequence. Hence, in most cases a *k*-mer index does not provide exact answers for queries of longer sequences (e.g., whole transcript or entire gene) but instead provides a reasonable approximation.

Data structures for indexing a set of *k*-mers have been studied in depth (Chikhi et al., 2019). Here, we consider the problem of storing a set of *k*-mer sets. A naive approach would be to use such an index separately for each *k*-mer set in the collection. However, a key aspect is that sequencing experiments that are analyzed collectively typically share a large fraction of *k*-mers. Therefore, significant space savings can be achieved by the identification and clever storage of this redundant information. Here, we focus on the methods underlying the different building blocks of sets of *k*-mer sets structures. We review the different properties, the types of queries, and the computational performance that they offer. We highlight similarities of methods based on commonalities between building blocks where it is appropriate.

## BIOLOGICAL APPLICATIONS

### RNA-seq studies

Transcriptomics was one of the first areas of application of the reviewed methods. Solomon and Kingsford (Solomon and Kingsford, 2016) gathered more than 2,500 samples of human RNA-seq, consisting of blood, brain, and breast tissue samples from SRA. This led to the possibility of identifying conditions which express isoforms by associating transcripts to tissues. Similarly to tissue-specific associations, one can envision the numerous benefits of comparing patient cohorts in order to understand differences in pathologies or impact of medication. For instance, using RNA-seq for functional alterations profiling has become more frequent in cancer research (Byron et al., 2016). Thus, vast programs such as The Cancer Genome Atlas (TCGA) (Tomczak et al., 2015), provide RNA-seq data from a variety of cancer types. Authors of SeqOthello (Yu et al., 2018) showed how to investigate gene-fusion using a set of *k*-mer sets by first creating an index of all tumor samples from the TCGA. Then they considered documented fusion events and their corresponding *k*-mer signatures, and screened the index to detect these signatures. They confirmed some fusions and reported some novel ones. Fusion transcripts provide interesting targets for cancer immunotherapies since they are prone to exhibit tumor-specific markers.

One of the data structures covered in this survey, the colored de Bruijn graph, has also been used for rapid, alignment-free quantification of RNA-seq data. Tools such as Sailfish (Patro et al., 2014), Salmon (Patro et al., 2017) and kallisto (Bray et al., 2016) rely on a colored de Bruijn graph (implemented using a hash table) to represent and quantify sets of transcripts per genes.

## Microbial genomics

Cortex (Iqbal et al., 2012), which introduced the concept of colored de Bruijn graph, was used to study the host diversity and dynamics of *Staphylococcus aureus* substrains using whole-genome sequencing (Young et al., 2012). Then, several papers demonstrated how sets of *k*-mer sets could be used to mine and analyze collections of microbial samples or genomes, whether they be strains of the same genera (e.g., 16,000 strain of *Salmonella)* using VARI-Merge (Muggli et al., 2019), microbiomes (e.g., 286,997 genomes from the human microbiome), or more extensive microbial data (e.g., 469,654 bacterial, viral and parasitic datasets from the ENA) using BIGSI (Bradley et al., 2019a). For example, GenomeTrakr (Timme et al., 2018) was developed to coordinate international efforts in sequencing whole genomes of foodborne pathogens. Indexing and querying this and other databases could lead to improved surveillance of pathogenic bacteria, and thus, elucidate the effectiveness of interventions that attempt to control them.

Subsequently, *k*-mer indices have been used to follow the spread of antimicrobial resistance (AMR) genes and plasmids across bacterial populations. The BIGSI authors also searched for plasmid sequences bearing AMR and initiated a study in an index containing a variety of microbial genomes. They identified some of these plasmids spread across different genera. Other AMR, such as SNPs associated with fluoroquine resistance, were studied across 100,000 *Salmonella* genomes with BiFrost (Luhmann et al., 2020).

Lastly, an effort was proposed to build a comprehensive human gut microbiome resource with the help of a set of *k*-mer set structure. Cultured genomes and metagenomes assembled from metagenomics data were combined in a BIGSI index to create the Unified Human Gastrointestinal Genome index (Almeida et al., 2020). This resource aims at exploration and enables looking for contigs sequences, genes, or genetic variants.

## Genome dynamics

In a study of fine-scale recombination landscape in birds, Cortex was used to *de novo* call variants in zebra finch raw datasets, bypassing the low-quality state of current genome resources (Singhal et al., 2015). Recently, a novel phylogeny approach was introduced by building a colored de Bruijn graph (Wittler, 2020) (using BiFrost) on a set of genomes (assembled or reads), and traverses the structure to extract phylogenetic signal. This approach bypasses the usual multiple genome alignment step.

As an aside, tools like Mash (Ondov et al., 2016) perform sketching of datasets, i.e. construct small sets of short *k*-mers that constitute signatures of datasets, in order to compute ecological distances between datasets. Since sketching uses specific techniques that do not rely on the entire *k*-mer sets (see a related review (Marçais et al., 2019)), we consider them outside the scope of this study.

Finally, beyond these applications, other topics are starting to be explored: integrated variant calling across large-scale gene, plasmid and transposon search (Blackwell et al., 2019, Bradley et al., 2019a, Miller et al., 2020), bacterial pan-genome indexation (Muggli et al., 2019), and gene fusion and pan-cancer analysis (Yu et al., 2018).

## QUERY MODEL

Here, we describe the types of queries that are supported by the surveyed methods.

Let 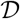 be a collection of *n* datasets. Let *S* be a nucleic sequence of arbitrary length such as a gene, or a transcript. Note that *S* can in principle be as short as a single *k*-mer, but in practice it is often a sequence longer than *k*. The aim is to determine the presence of *S* in each dataset of 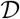.

The most elementary type of queries supported by all methods in this survey consists in reporting every dataset 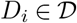 which contains a query *k*-mer. It can be naturally extended to a longer query sequence *S* by querying each element of the set *Q* of all *k*-mers present in *S*. One can see that if *S* is present in a dataset, then every element of *Q* is present as well. However the converse is not true: *k*-mers in *Q* may possibly correspond to different sequences within a dataset and *S* may actually be absent. Consider the two *k*-mers ACT and CTG (*k* = 3) and assume they are present in two different reads within a single dataset. A query sequence ACTG would then be reported as present in that dataset, regardless of whether the sequence is truly found as part of a single read. Despite this potential shortcoming, this is widely considered to be a reasonable approximation, due to *k* being long enough to make such false positive events unlikely.

In a seminal work, Solomon and Kingsford (Solomon and Kingsford, 2016) proposed to report every dataset 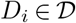 in which a proportion of at least *θ k*-mers of *Q* appear, i.e. |*D_i_* ⋂ *Q*|*/*|*Q*| ≥ *θ.* Here *θ* can be seen as a stringency parameter for the search. This query model is motivated by events such as sequencing errors and variants, which reduce the number of common *k*-mers between a query and a target. Thus, it is interesting and often necessary to report when only a fraction of the *k*-mers from a query sequence are present a dataset. Typically *θ* is set between 0.7 and 0.9 (Solomon and Kingsford, 2016). Also, the typical *k*-mer size range seen in applications is 21–31.

## BUILDING BLOCKS

We view the storage of a set of *k*-mer sets as having four possible components. These components are: the underlying data structure used to represent a single set, the strategy used to aggregate *k*-mers across different sets, the data structure used to store this aggregation, and the compression strategy used. See Figure 2 for an illustration of this view. Most of the novelty in the methods comes in the aggregation strategy and in the data structure used to support it. The other two components are used more as building blocks. In this section, we will cover these two building blocks.

**Fig. 2.**
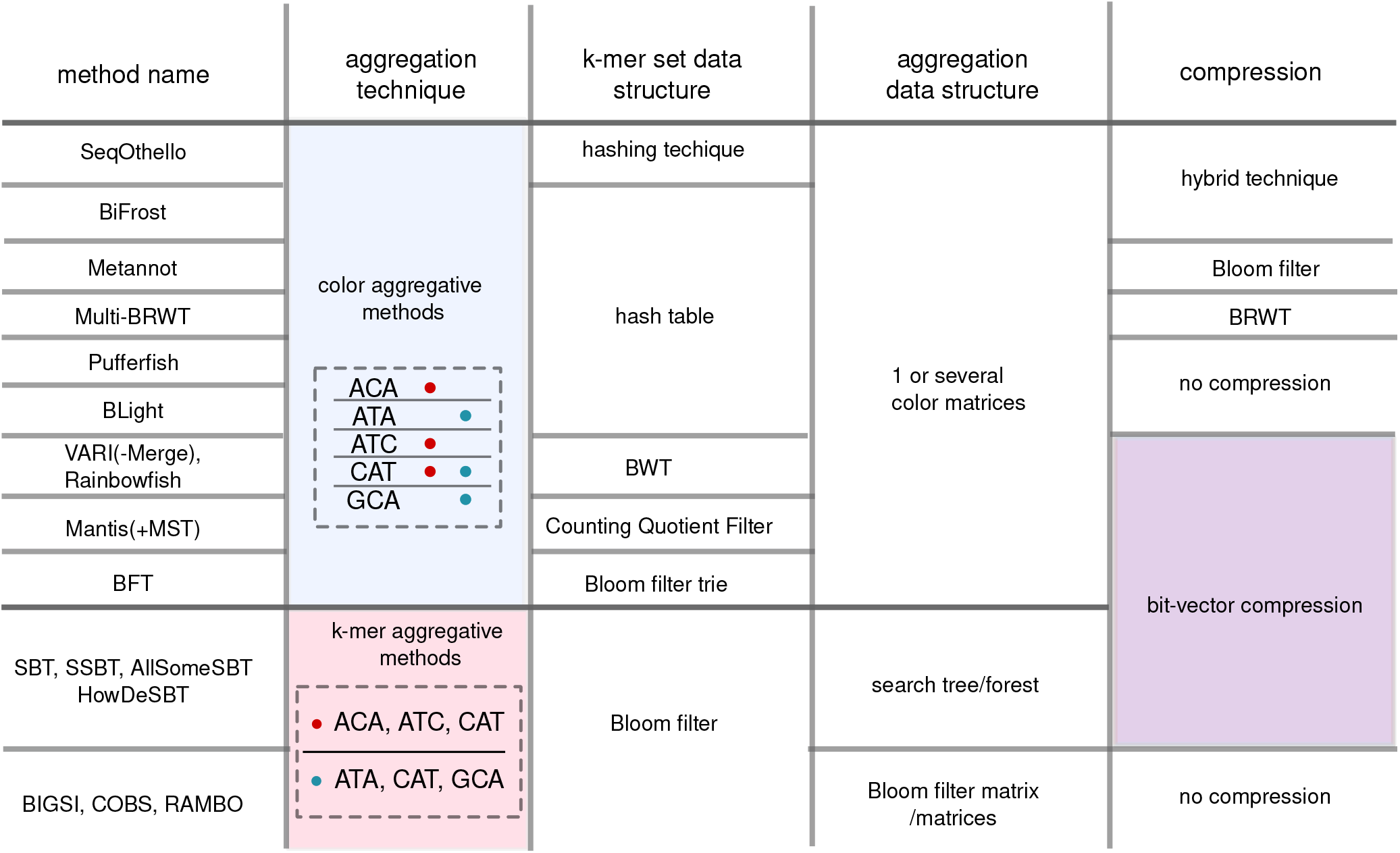
Overview of set of *k*-mer sets building blocks. We classified strategies in color-aggregative approaches, and *k*-mer aggregative approaches (second column). The top row of the figure indicates the general categories of components of each methods: the type of *k*-mer set; the way multiple sets are combined together; and an optional compression scheme. Each next row describes one of the surveyed methods. The cells in this figure are methodological choice, potentially common across methods, hence many cells are joined.

### *k*-mer set data structures

A set of *k*-mers is a collection of *k*-mers that are de-duplicated and unordered. It will be enough to consider them as black boxes that support all or a subset of the following operations:

- <u>membership</u>, i.e., testing whether a *k*-mer is in the set;
- <u>insertion</u> and/or <u>deletion</u> of a *k*-mer.

However, methods to represent *k*-mer sets are not all equivalent in terms of features and performance. We briefly review their main characteristics in the rest of this section but refer the reader to the recent survey by Chikhi et al. (Chikhi et al., 2019) for further details.

Most methods rely on bit vectors to store the presence or absence of *k*-mers in datasets. A bit vector is an array of bits, e.g., 00101 represents a bit vector of length 5. A 0 is used to indicate that the *k*-mer is absent, and a 1 indicates that it is present. A bit vector could be used to record the datasets in which a given *k*-mer appears, or, alternatively, all the *k*-mers that are contained in a given dataset. However, with a growing number of datasets and *k*-mers, using plain bit vectors is generally too simplistic, and often compression or other tricks are also incorporated. One example is the Bloom filter (Bloom, 1970), which is a way to store a set as a bit vector using many fewer bits than the naive approach (see Box 1).

##### Box 1. Technical definitions

<u>Hashing, hash functions</u>. Mathematical functions that are used to associate elements (e.g. *k*-mers) to numbers (e.g. positions in an array).

<u>Bloom filters</u>. Bit vectors that record the presence or absence of elements within a set, with some approximation, using hashing.

<u>Counting Quotient filters (CQFs)</u>. Similar in nature to Bloom filters but differ by their hashing strategy. CQFs support membership queries of elements in a set, and also counting elements in a multi-set.

<u>Minimal perfect hash functions (MPHFs)</u>. Functions that associate a fixed set of elements to the range of consecutive integers from 0 to the number of elements, in a highly space-efficient way.

See Supplemental Box S1 for detailed illustrations of BF, CQF and a hashing method inspired by MPHF techniques called Othello.

<u>Burrows Wheeler transform (BWT)</u>. A text transformation algorithm. Given an arbitrary text, such as a DNA sequence, BWT rearranges it in a way that enhances its compression and permits indexing. The transformation is reversible, allowing the text to be efficiently recovered.

###### Graph definitions

A <u>graph</u> (see (a) in figure below) is a pair of two sets *V* and *E*. Elements of *V* are nodes (in blue), and elements of *E* are pairs of related nodes called edges (in orange and red).

A <u>path</u> in a graph is a sequence of edges that connects a sequence of distinct nodes (the example shows a path drawn between nodes 1,2,3,4 through red edges).

A <u>tree</u> ((b) in figure below) is a particular graph in which any two nodes are connected by exactly one path. A forest is a disjoint union of trees ((c) in figure below). A <u>subtree</u> (circled in grey in (b)) is a subset *G^′^* and *E^′^* of a tree *T* = (*G,E*).

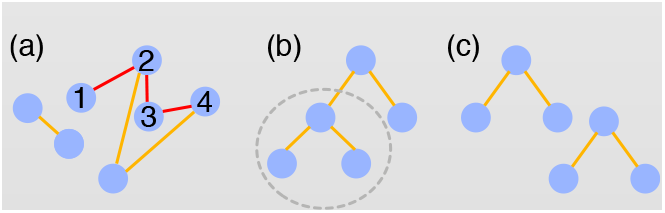

A <u>trie</u> is a tree that allows to efficiently store and query a set of words. Wavelet tries (Grossi and Ottaviano, 2012) are designed to store compressed sequences.

A <u>de Bruijn graph (DBG)</u> is a graph where nodes are *k*-mers and there exists a directed edge from vertex *u* to *v* if the last *k* – 1 characters of *u* are the same as the first *k* – 1 characters of *v*. <u>A compacted de Bruijn graph</u> is a different graph than the DBG, which however represents the same *k*-mer information by merging unambiguous paths.

See Supplemental Box S2 for an example of DBG and compacted DBG.

Some methods view a *k*-mer set as a de Bruijn graph (DBG, see Box 1). These two views, *k*-mer sets and DBGs, are (in some sense) equivalent as they intrinsically represent the same information.

Data structures for representing *k*-mer sets (DBG or not) can further be divided into two categories: membership data structures and associative data structures. The first category only informs about the presence or absence of *k*-mers, e.g., as in the case of a Bloom filter. The second category associates pieces of information to *k*-mers, akin to how dictionaries link words to their definitions. Some examples of associative data structures include hash tables and counting quotient filters (CQF) (Almodaresi et al., 2019, Bender et al., 2012, Pandey et al., 2017) (Box 1).

Some data structures (e.g. CQF, BOSS) can represent sets in an exact way; whereas others (e.g. Bloom filters) represent them in a probabilistic way, meaning that the structures can return false positives (i.e., meaning that a *k*-mer is sometimes falsely reported as present in the set when in fact it is absent). These false positives lead to an over-estimation in the number of *k*-mers detected as present in a set. While this is an undesirable effect, it can be partly mitigated – as we will see in Section Background and method intuition.

The main reason such advanced data structures are considered, instead of those provided in the standard libraries of programming languages, is space efficiency. Bloom filters and CQFs approximately require a byte for each element in the set, i.e., less than the size of the element itself. Similarly, optimized representations of DBGs (Boucher et al., 2015, Bowe et al., 2012, Chikhi and Rizk, 2013), which are also representations of *k*-mer sets, aim for near-optimal space efficiency. Exact and probabilistic data structures offer a trade-off between space and accuracy. This is a crucial aspect as the volume of data typically exceeds what can be processed using unoptimized data structures.

### Compression

To further optimize space usage, different compression techniques have been applied to sets of *k*-mer sets. Bloom filters and bit vectors are amenable to a number of compression techniques because they are represented in bits. They can be sparse (i.e. when most of the bits are 0s) or dense (i.e. when most of the bits are 1s). Compression methods exploit these properties.

### Bit vector compression

Bit vectors can be efficiently stored using bit-encoding techniques that exploit their sparseness or redundancy. The most prevalent of the methods in this survey are RRR (Raman et al., 2002) or Elias-Fano (EF) (Elias, 1974, Fano, 1971, Ottaviano and Venturini, 2014). The principle behind these is to find runs of 0’s and to encode them in a more efficient manner, reducing the size of the original vectors. Other techniques such as Roaring bitmaps (Lemire et al., 2016) adjust different strategies to sub-parts of the vectors. Wavelet trees (Grossi et al., 2003) generalized compression of vectors on larger alphabets (i.e., not just 0’s and 1’s but e.g. a’s, b’s, c’s, etc). More advanced techniques deriving from the same concept were also proposed specifically for sets of *k*-mer sets (Karasikov et al., 2019).

### Delta-based encoding

When two sets share many elements, it may be more advantageous to store only one along with the differences with the other. For instance, rather than storing two (possibly compressed) bit vectors, one can only store the first bit vector explicitly along with a list of positions that need to be inverted to obtain the second vector.

This list of positions can itself be encoded as a bit vector, with a bit set to one if and only if the bit is at the location of a difference between the two vectors. This bit vector is expected to be more sparse than the original encoding, allowing better compression. Such a scheme is usually called “delta-based encoding” in the literature.

### Hybrid techniques

In hybrid approaches, a collection of bit vectors is split into different buckets, where each bucket contains bit vectors with similar features. Compression within each bucket is performed using a suitable technique selected from a pool of feature-specific ones. An illustration of these techniques is provided in Supplemental Box S4.

## COLOR-AGGREGATIVE METHODS

The methods that we will survey are split into two categories. Color-aggregative methods index the <u>union set</u> of *k*-mers, which is the joint set of all *k*-mers that appear in all the datasets. Instead, *k*-mer aggregative methods index each dataset separately, then build an aggregation data structure to distribute queries. A few methods that escape this categorization will be presented separately.

Color-aggregative methods gather and index the union set of *k*-mers, then associate information to each *k*-mer to record its dataset(s) of origin. A practical advantage of color-aggregative methods is that a *k*-mer that appears in many samples will appear only once in the union set. This greatly reduces redundancy in the representation of *k*-mers, but introduces the need to store additional color information. In this subsection, we give a brief background and history of the methods that fall into this category.

### Color matrix

A <u>color</u> of a given *k*-mer is frequently used in the literature to identify a dataset containing the *k*-mer, assuming each dataset is given a unique color. A <u>color set</u> is the set of colors associated with a *k*-mer. It is convenient to represent a color set using a bit vector. Here, fixing an ordering of the datasets, a 1 at position *i* in the bit vector indicates the presence of the *k*-mer in the *i*-th dataset, and 0 its absence. Given *n k*-mers and *c* datasets, the <u>color matrix</u> is a *n* × *c* matrix of bits which describe the presence or absence of each *k*-mer (in the rows) in each dataset (in the columns). For an example, see Box 2.

##### Box 2. Representation of colors in color-aggregative methods

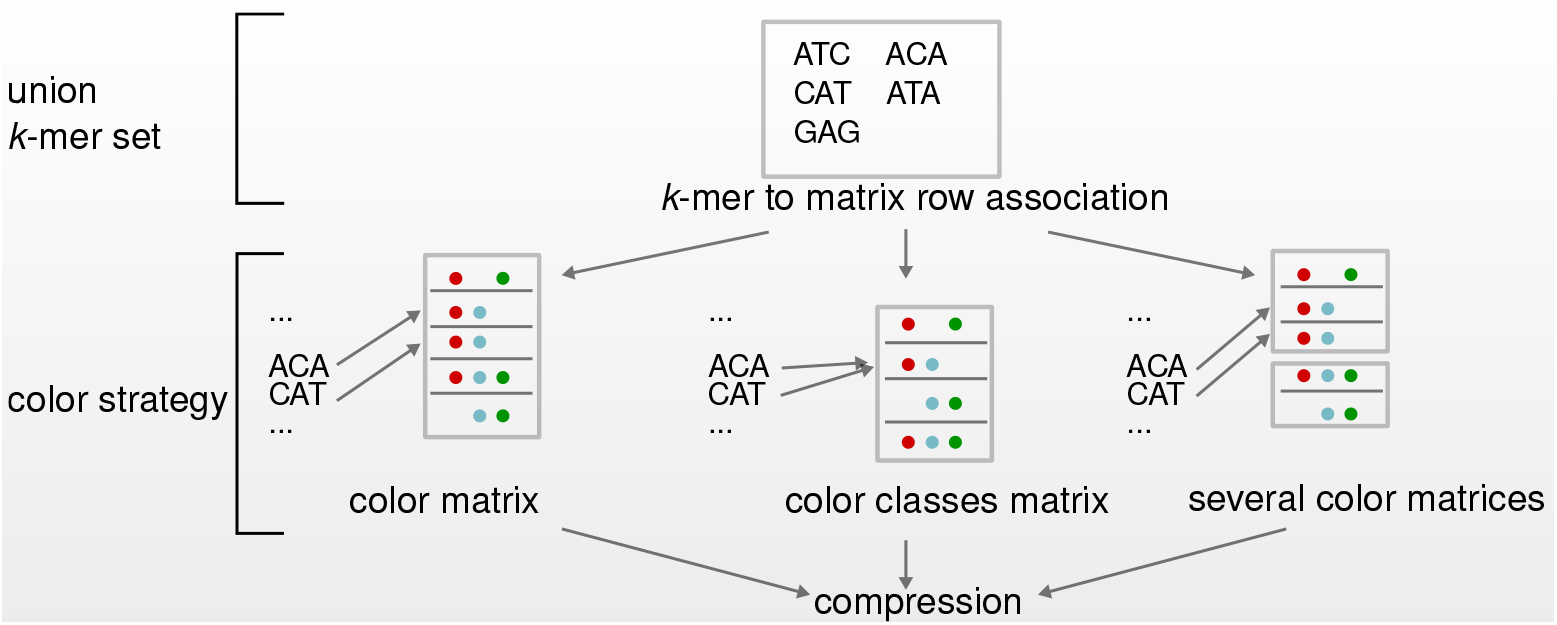

A **color matrix** represents the presence of *n k*-mers across *c* datasets (in the above figure, *n* = 5*, c* = 3). Different schemes have been introduced to encode such matrices. In particular, a **color class** is a set of colors that is common to one or more *k*-mers. In other words, in the color matrix there may be identical rows, then the corresponding *k*-mers belong to the same color class. One may “de-duplicate” the *n* rows of the color matrix into *m* < *n* color classes (here, *m =* 4). Then, color class identifiers are introduced as intermediaries between *k*-mers and color classes. (To go further, frequently used color classes can be referenced using fewer bits by using small integers as identifiers).

###### Color-aggregative methods

are generally composed of three parts. First, a set of all *k*-mers, built using either a DBG or an ad-hoc representation. Second, a correspondence between *k*-mers and colors in the form of a color matrix (left, ‘color strategy’ row), color classes (middle), or several color matrices (right). Finally, *k*-mer sets and/or colors may be further compressed. In the two first columns’ strategies, compression is based on one of the techniques from section Compression. In the third column, different techniques may be used for different matrices.

### Background

The first color-aggregative method was proposed by Iqbal *et al.* (Iqbal et al., 2012) in the Cortex software. It implements a relatively straightforward associative data structure that maps *k*-mers to colors. A <u>colored DBG</u> is a DBG of the union set of *k*-mers, where each vertex is labelled with the color set of the corresponding *k*-mer. Iqbal *et al.* used a colored DBG built on a set of individuals from a population to detect SNPs and other short variants; such events are reflected in the graph by a pair of short paths that share their start and end nodes. This enabled to detect genetic variation in a population without the use of a reference genome. Cortex consumes an inordinate amount of RAM when the total number of distinct *k*-mers exceeds tens of billions. This main drawback motivated more recent works improving the efficiency of the colored DBG.

We note that later in this survey, we will not restrain the term colored DBG to the original work from Iqbal et al., but will extend it to any explicit DBG implementation that associates color sets to *k*-mers. Colored DBGs are only implemented in the class of color-aggregative methods. Conversely, as we will see later, some color-aggregative methods do not implement a colored DBG.

### Later methods

The first color-aggregative methods that improved upon the memory- and time-efficiency of Cortex were Bloom filter trie (BFT) (Holley et al., 2016) and Vari (Muggli et al., 2017). These methods achieved a significant reduction in representation size via different strategies. After the introduction of these methods, subsequent improvements were made with the development of Rainbowfish (Almodaresi et al., 2017), Multi BRWT (Karasikov et al., 2019), Mantis (Pandey et al., 2018), SeqOthello (Yu et al., 2018), Mantis+MST (Almodaresi et al., 2019), and Vari-Merge (Muggli et al., 2019). Most of these recent techniques rely on a more careful encoding of the colors of each *k*-mer, which takes advantage of redundancy in the data.

A summary of the main features of color-aggregative methods is presented in Table 1. Their methodological strategies are presented in more details in Supplemental Box S6.

**Table 1.** Summary of the existing color-aggregative methods and some of their features. **nav.**: indicates if it is possible to navigate in the DBG (ie. going from one *k*-mer to the following ones and conversely). Such a navigation allows for instance to perform variant-calling. We note that these two aspects should be technically possible in all colored DBG tools. **add**: indicates if new datasets can be added to the index. Although it is conceptually possible that new data can be added to Vari, Rainbowfish (by rebuilding) and Mantis, this feature is currently not implemented. **exact**: indicates if the index provides exact results (Y) or if there may have false positives (N).

| Method | Description | Nav. | Add | Exact | Remarks |
| --- | --- | --- | --- | --- | --- |
| VARI | A BWT-based index on $k$-mers interfaced with a color matrix compressed with RRR | Y | N | Y | Supports short variant discovery |
| VARI-Merge | A divide-and-conquer approach to building VARI on large datasets | Y | Y | Y |  |
| BFT | A tree that records $k$-mers, using Bloom filters, and their compressed color classes | Y | N | Y | First scalable color-aggregative method |
| Rainbowfish | Similar to VARI for the $k$-mers, similar to BFT for the colors | Y | N | Y | |
| Mantis | A tweaked CQF that records $k$-mers and color classes, which are compressed with RRR | Y | N | Y/N | Can switch to probabilistic queries to save space |
| Mantis+MST | Similar to Mantis with more efficient delta-encoding color compression |  |  |  |  |
| SeqOthello | MPHF (Othello) that maps $k$-mers to their color sets grouped and compressed by similarity using hybrid bit-vector compression | N | N | N | Most memory efficient among color-aggregative methods |
| Pufferfish, BLight | A MPHF-based hash table associating $k$-mers (in a DBG) to colors | N | N | Y | |
| BiFrost | A hash table associating $k$-mers to several color matrices (similarly to Mantis+MST) with adapted compression strategy | Y | Y | Y | |
| Metannot | A hash table storing $k$-mers and nearly optimal compressed colors with wavelet tries and RRR | N | Y | Y | Can delete datasets |
| Multi-BRWT | A hash table storing $k$-mers with a color matrix compressed in both dimensions simultaneously | N | Y | Y | Can delete datasets |
| REINDEER | A MPHF-based hash table associating $k$-mers to counts | N | N | Y | Only method that allows to store counts |

### Methods based on a color matrix

The representation of the DBG used by **Vari** (Muggli et al., 2017) is a BWT implementation referred to as BOSS (Boucher et al., 2015), While we will not go into the technical details here (see Supplemental files and Supplemental Box S3 for more information), it is sufficient to see BOSS as a rearrangement of the original data that enables indexing and compression. In order to add color information in Vari, a compressed color matrix is constructed row-by-row.

Later, **Vari-Merge** (Muggli et al., 2019) was introduced to construct colored DBGs for very large datasets, which can also be updated with new data. Two other methods, **Pufferfish** (Almodaresi et al., 2018) and **BLight** (Marchet et al., 2020b) put emphasis on the *k*-mer indexing technique in order to efficiently store DBGs in memory, and use a simple color matrix to represent the colored DBG. It is worth noting that BLight shares similarities with Kraken (Wood and Salzberg, 2014), a taxonomic classifier. Indeed, it can be seen as a colored de Bruijn graph of *k*-mers with labels to their genome of origin.

### Methods based on color classes

In many applications, such as human RNA-seq indexing (Solomon and Kingsford, 2016), it is expected that many datasets share a large number of *k*-mers. This redundancy can be exploited to reduce the color encoding size, through <u>color classes</u>. When colors are seen as bit vectors, the color classes are defined simply as de-duplicated bit vectors. Thus two *k*-mers having the same color sets are associated with a single color class instead of two identical bit vectors (see Box 2). Compression may be achieved by representing the color matrix as a compressed bit vector.

**Bloom Filter Trie** (Holley et al., 2016) (BFT) uses a different approach to storing the DBG than Vari but also aims at representing a DBG. BFT introduced the idea of color classes, and a more detailed description of its inner workings is provided in Supplemental Box S5. **Rainbowfish** (Almodaresi et al., 2017) mixes ideas from Vari and BFT. **Mantis** (Pandey et al., 2018) introduces another strategy (the CQF, see Box 1) for storing the DBG in a space-efficient manner. Initially, CQFs were introduced to record counts associated with *k*-mers but in Mantis, the structure instead stores color sets. **Mantis+MST** (Almodaresi et al., 2019), an extension of Mantis, takes advantage of the insight that many color classes are frequently similar to each other, since many *k*-mers occur in relatively similar sets of sequences. Thus it proposes a more efficient encoding of colors.

### Methods based on several matrices

**SeqOthello** (Yu et al., 2018) does not explicitly represent a DBG but rather stores a probabilistic set of *k*-mers using a hashing method inspired by MPHF techniques called Othello (see Supplemental Box S1). SeqOthello proposes to group similar color profiles, then uses a suitable compression technique depending on the sparsity of each group. It may wrongly associate a dataset to an alien *k*-mer, instead of correctly returning that such *k*-mer belongs to no dataset. For a query that consists of many *k*-mers (such as genes or transcripts) errors can be mitigated because false positives are unlikely to all point to the same dataset(s). This property will also be used in *k*-mer aggregative structures, and will be further discussed in Section Background and method intuition.

Recently, another construction strategy for color-aggregation was proposed in **Bifrost** (Holley and Melsted, 2019). As in BLight and Pufferfish, BiFrost uses a <u>compacted DBG</u> (introduced in e.g. (Chikhi et al., 2016, Minkin et al., 2016), see Supplemental Box S2), to more efficiently represent the sequences than a set of individual *k*-mers. In addition, we note these methods are similar to Mantis and SeqOthello in terms of the underlying hash-based strategies, and the differences are detailed in Supplemental Box S6.

### Other methods

Some techniques evade the above categorization and have focused on specific aspects of color-aggregative methods memory optimization. The growing number of colors (or classes) motivated works to further reduce space through <u>lossy compression</u>. In **Metannot** (Mustafa et al., 2018) and **Multi-BRWT** (Karasikov et al., 2019) the main contribution is not the data structure used to store the graph, but the one used to store colors. Metannot explores two strategies for color compression. One of them is probabilistic: in order to reduce false positives, color sets queries are corrected by taking the intersection with other color sets from neighboring *k*-mers in the DBG. Multi-BRWT improves upon standard bit-encoding representation (such as RRR, EF).

Colored DBGs have been used to perform RNA-seq quantification (Bray et al., 2016, Patro et al., 2017, 2014), by associating colors to individual genes as opposed to datasets. Yet such methods still require a pseudo-alignment step to recover abundance information from the reads. Recently, **REINDEER** (Marchet et al., 2020a) proposed a color-aggregative index which also records abundance, bypassing the need to align reads in order to recover abundances. It relies on BLight, to which it adds novel features in indexing and a more advanced color matrix with color classes and compression.

### Queries

Given that current implementations use *k*-mers that fit within (extended) computer words (~ 21 to 63), the query time bottleneck comes mainly from random memory accesses, neglecting the time taken to calculate and hash the *k*-mers. Hash-based methods perform very fast <u>color</u> queries: retrieving <u>information relative to a single</u> *k*-mer requires only a constant number of memory accesses. The methods whose underlying DBG is BOSS (e.g., Vari, Rainbowfish, Vari-Merge) are expected to show a lower throughput. Indeed, the retrieval of a *k*-mer requires in the order of *k* memory accesses.

## *K*-MER AGGREGATIVE METHODS

We now turn to a completely different class of data structures. Unlike previously mentioned methods, *k*-mer aggregative methods do not pool *k*-mers from all datasets in order to build an index. Rather, they first process datasets separately, and then aggregate them in different ways. A summary of these methods’ features appears in Table 2.

**Table 2.** Summary of the existing *k*-mer aggregative methods and some of their features. **Add**: indicates if new datasets can be added to the index. * These methods’ papers propose a dataset addition algorithm though it is not implemented. **exact**: indicates if the index provides exact results (Y) or if there may have false positives (N). **PRQ**, stands for Partial RAM query: indicates if the query can be performed by loading only a small part of the index in RAM. This is much less space-consuming but potentially less time-efficient. Comparatively, all color-aggregative methods need to load the whole index in memory when querying. However, contrarily to some color-aggregative methods, no *k*-mer aggregative method offer navigation operations.

| Method | Description | Add | Exact | PRQ | Remarks |
| --- | --- | --- | --- | --- | --- |
| SBT | A tree of RRR-compressed BF's with each dataset stored in a leaf | N | N | Y |  |
| SSBT | A tree similar to SBT but with more fine-grained BF's. | N* | N | Y | Redundancy removal compared to SBT |
| AllSomeSBT | A tree similar to SSBT but with a hierarchical clustering of datasets to save space and query/construction time | N* | N | Y |  |
| HowDeSBT | Similar to AllSomeSBT but with additional BF optimizations | N | N | Y | Small index |
| BIGSI | A matrix of BF's, where each column corresponds to a dataset | Y | N | N |  |
| COBS | Similar to BIGSI with BF's of variable lengths to adapt for varying dataset sizes | N | N | Y | Fast construction |
| RAMBO | A matrix of BF's, where each BF corresponds to an union of several datasets | Y | N | N |  |

### Background and method intuition

All the *k*-mer-aggregative methods surveyed work by storing the *k*-mers of each dataset in a separate Bloom filter (BF), i.e. using one BF per dataset. A BF is a probabilistic data structure that sometimes returns false positives; i.e. the BF may report that a *k*-mer belongs to a certain dataset when it really does not. In the query model that we defined on Section Query model, the *θ* parameter allows to partially mitigate this problem by considering the results of multiple *k*-mer queries. Indeed, the false positive rate for a sequence decreases exponentially with the number of *k*-mer queries (Bingmann et al., 2019, Solomon and Kingsford, 2016). For the values of *θ* used in practice, the false positive rate of an individual BF can be set as high as 50% without degradation of query performance on sequences that are much longer than *k* (Solomon and Kingsford, 2016).

Prior to construction, unlike color-aggregative methods, most *k*-mer-aggregative methods (except COBS) need to know in advance how large is the largest dataset, in terms of the number of distinct *k*-mers, to ensure that BFs are appropriately sized. The common BF size is chosen according to a desired false positive rate.

### *k*-mer aggregative methods summary

#### Sequence Bloom trees (SBTs)

These are a family of related techniques detailed across multiple publications (Harris and Medvedev, 2020, Solomon and Kingsford, 2016, 2018, Sun et al., 2017), adapted to bioinformatics from a previously-known hierarchical structure of BFs (Bloofi (Crainiceanu and Lemire, 2015)). The tree structure represents a hierarchical clustering of the datasets, e.g. one obtained using *k*-mer similarity across datasets. In the original SBT (Solomon and Kingsford, 2016), each leaf corresponds to an individual dataset and each internal node is a BF that represents all the *k*-mers of the datasets descendant from it. Split-Sequence Bloom trees (**SSBT**, (Solomon and Kingsford, 2018)) and **AllSomeSBT** (Sun et al., 2017) simultaneously proposed to instead store two BFs per internal node, each one separately containing the *k*-mers present in (respectively. absent from) all the descendants. **HowDeSBT** (Harris and Medvedev, 2020) further improved the space utilization and provided the first analytical analysis of the running time and memory usage of the various SBT approaches. These recent improvements (Harris and Medvedev, 2020, Solomon and Kingsford, 2018, Sun et al., 2017) greatly reduced the space and query time compared to the original SBT (see Supplemental Box S8 for a comparison of SBT flavors).

#### Matrix strategies

An orthogonal approach was proposed in **BIGSI** (Bradley et al., 2019a) (Box 3 (b)). As a first approximation, BIGSI can be seen as the concatenation of many BFs, forming a color matrix. The matrix is stored in row-major order, i.e. row-by-row, so that each row appears as a consecutive block and can be efficiently queried. A closely related work is presented as a part of DREAM-Yara (Dadi et al., 2018), where BFs are interleaved, in order to efficiently retrieve the same position of several BFs (see Supplemental Box S8). BIGSI was later improved, speed and memory-wise, by COBS (Bingmann et al., 2019).

###### Data structures

In *k*-mer aggregative strategies, Bloom filters representing each of the datasets are organized in either a tree or a matrix structure. In the example below, there are four datasets (red, blue, green, and yellow), and the grey rectangles represent Bloom filters.

a. **Tree strategy** A search tree is constructed, where each leaf is a dataset and internal nodes represent groups of datasets. Datasets with similar BFs can be clustered to reside in the same subtree. In the original SBT approach (Solomon and Kingsford, 2016), each node stores exactly one BF, containing all the *k*-mers present in the datasets of its subtree. For a leaf, this is simply the *k*-mers in the corresponding dataset. Later versions of SBTs (Harris and Medvedev, 2020, Solomon and Kingsford, 2018, Sun et al., 2017) store more sophisticated data at each node, though they still rely on BFs.
b. **Matrix strategy** The BFs from all the datasets are concatenated column-wise to obtain a matrix similar to a color matrix. A row in the matrix roughly corresponds to a *k*-mer (more precisely, to the position indicated by the hash value of a *k*-mer). In the original BIGSI approach (Bradley et al., 2019a), all BFs have exactly the same size. In COBS (Bingmann et al., 2019), datasets of comparable cardinality are grouped into bins, leading to a collection of matrices of different sizes.

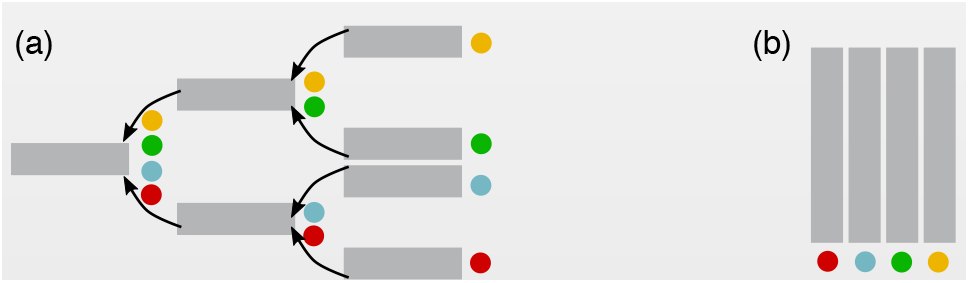

###### Queries

Consider an example query composed of 3 *k*-mers with a threshold 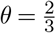 where, for simplicity, each *k*-mer corresponds to a single location in the BF.

a. **Tree strategy** Conceptually, one starts at the root node and then explores down the tree, always checking all the children of a node before moving to another node (breadth-first strategy). A counter of *k*-mers that have been determined to be present or absent is maintained for the query as it propagates down the tree. If either of the counters exceeds a certain threshold, the search does not propagate further down the subtree of that node. E.g., in panel (a) below, black bars in the BF represent the presence of 3 queried *k*-mers. Considering that 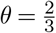 of the *k*-mers should be present, the query is pruned at the yellow/green nodes since not enough present *k*-mers are found. Only the red dataset is returned as containing the query.
b. **Matrix Strategy** In BIGSI, each *k*-mer corresponds to a single row in the matrix which is then extracted, and summed column by column to obtain a vector where each element contains the number of *k*-mers occurring in the corresponding dataset. Again in panel (b) below, black bars in the BF represent the presence of 3 queried *k*-mers. The red dataset will be the only one returned as containing the query.

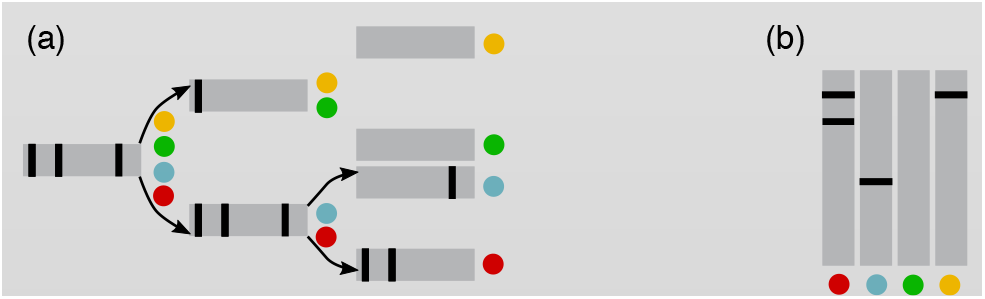

Detailed examples can be found in Supplemental Box S7. **RAMBO** (Yan et al., 2019) appears at first glance related to BIGSI, as it is also a matrix of BFs. However, RAMBO is actually closer to SBTs: each BF in the RAMBO matrix represents several datasets, but not in a hierarchical fashion. More details are provided in the Supplemental files and Supplemental Box S8.

### Queries

In SBTs, a query containing several *k*-mers *Q* starts at the root and propagates down the tree. At any node, the information stored at that node is used to determine which *k*-mers of *Q* are determined in all descendants of the node. Depending on the SBT flavor, a determined *k*-mer may be for sure present, or for sure absent, in all descendants. The *k*-mers which are determined are discarded from *Q* as the query propagates down the tree (see Box 3). When enough *k*-mers become determined, the search can be pruned, i.e. not carried further down the tree.

In matrix approaches, a *k*-mer query extracts a bit vector indicating its presence across D (see Box 3). For a set of *k*-mers *Q*, the bit vector for each *k*-mer in *Q* is constructed and bitwise operations on these vectors are performed to answer the whole query.

### Other schemes

Other unpublished tools have considered different techniques for storings and indexing sets of *k*-mer sets. **kamix** (https://github.com/jaudoux/kamix) uses SAMtools’s BGZF library (block-compressed gzip) to store and index a *k*-mer matrix. From the same author, **kad** (https://github.com/jaudoux/kad) uses a RocksDB database to store a list of *k*-mers and counts.

BEETL (Cox et al., 2012) is a technique that stores inside a BWT all sequences (i.e., not *k*-mers, but the original data) from a sequencing dataset. BEETL was able to compress and index 135 GB of raw sequencing data into a 8.2 GB space (on disk for storage, or in memory for queries). A variant, BEETL-fastq (Janin et al., 2014), also enabled to perform efficient sequences searches and was also applied to the representation of multiple datasets.

Population BWT (Dolle et al., 2017) is also a scheme based on BWT geared towards the indexing of thousands of raw sequencing datasets. The BWT allows to query *k*-mers of any length and additionally gives access to the position of each *k*-mer occurrence within the original reads (note however that reads need to be error-corrected).

Recently, compressed structures able to compress full-text were proposed as proofs of concept for indexing and querying collections of biological datasets (Cobas et al., 2020) for presence/absence and abundance. However, at the time of writing, such indexes have been tested on a few dozens of close bacterial strains, not yet on raw reads data.

## PERFORMANCES OVERVIEW

### Index construction on human RNA-seq samples

Indexing datasets of a similar type, such as RNA-seq samples from the same species, was one of the first applications proposed in the literature of sets of *k*-mer sets, and remains one of the main benchmarks for these tools. Table 3, based on Supplemental Tables S1 and S2, reports the performance of most of the recent tools on a collection of human RNA-seq datasets (2,652 RNA-seqs from the original SBT article (https://www.cs.cmu.edu/~ckingsf/software/bloomtree/srr-list.txt). This table was constructed by gathering results from three recent articles (Bradley et al., 2019a, Harris and Medvedev, 2020, Yu et al., 2018). As the articles use different hardware and slightly different parameters, a direct comparison of the tools is challenging. Instead, Table 3 presents a summary of the best possible performance that can be currently achieved on the given datasets. Supplemental Table S3 also presents a summary of methods’ time complexities.

**Table 3.** Overview of the best achievable performance in terms of space and time requirements to build indices. The data processing time column refers to the time necessary to convert the original sequence files to the *k*-mer set indices (Bloom filters / Othello). The maximum external memory column corresponds to the peak disk usage when building the index. The time column is the time required to build the set of *k*-mer sets index (on one processor). The index size column is the final index size.

| Data set | Data Processing Time (days) | Max Ext. Memory (GB) | Time (h, wallclock) | Peak RAM (GB) | Index Size (GB) |
| --- | --- | --- | --- | --- | --- |
| Human RNA-seq<br>(2,652 datasets) | 2.5 (Harris and Medvedev, 2020)<br>(HowDeSBT) | 30 (Harris and Medvedev, 2020)<br>(HowDeSBT) | 2 (Yu et al., 2018)<br>(SeqOthello) | 5 (Yu et al., 2018)<br>(SSBT) | 15 (Harris and Medvedev, 2020)<br>(HowDeSBT) |
| Bacterial genomes<br>(1,000 datasets) | Not reported | Not reported | 0.01 (Bingmann et al., 2019)<br>(COBS) | 1.5 (Bingmann et al., 2019)<br>(SSBT) | 1.9 (Bingmann et al., 2019)<br>(HowDeSBT) |

The data processing phase, where the *k*-mer sets are constructed and initialized, often takes significant time across all methods. It is usually not presented as a bottleneck since it is viewed that this step can be computed while downloading the samples.

Regarding query times, each method has used different experimental setups, making comparisons difficult, e.g., using transcript batches of different sizes (100-10,000). We refer the reader to the experimental benchmark in (Harris and Medvedev, 2020), which compares the average query times for randomly selected batches, effects of warming the cache, maximum peak memory for queries. Finally, we note that the information output by a query can vary from one implementation to another and that the maximum supported value for *k*-mer size is also implementation dependent.

### Indexing bacterial genomes

We now turn to the indexing of large collections of bacterial datasets. Table 3 summarizes the benchmark published in the COBS article (Bingmann et al., 2019). Methods that were reported to perform the best in the previous section remain the most efficient, and most recent methods tend to show the best performance. BIGSI, AllSomeSBT, and COBS queries are fast. We note that COBS construction time is very low (< 1 minute), in part because it has been run using 80 threads. SSBT and COBS also have low RAM consumption. An advantage of HowDeSBT is the small size of the index on disk. This demonstrates that highly-diverse datasets, in terms of *k*-mer contents, can also be efficiently stored in variants of SBTs.

### Indexing human genome sequencing data

To the best of our knowledge, only two methods (BEETL-Fastq and Population BWT) have been applied to the representation of full read information from cohorts of whole human genomes. BEETL-Fastq represented 6.6 TB of human reads in FASTQ format in 1.7 TB of indexed files. Population BWT managed to index (in a lossy way) 87 Tbp of data, corresponding to 922 billion reads from the 1,000 Genomes Project. After read correction and trimming, the authors obtained a set of 53 billion distinct reads (4.9 Tbp) and indexed it with a BWT stored with 464 GB on disk (requiring 561 GB of main memory for query). Metadata (e.g., sample information for each read) was stored in a 4.75 TB database.

Given the apparent difficulty to construct large-scale indices on human sequencing datasets, we conclude that this is not yet a mature operation. Therefore, we did not provide a detailed comparison with other indexing techniques.

## DISCUSSION

General observations can be derived from the comparison we have presented in this survey.

SBT approaches were designed for collections with high *k*-mer redundancy, such as human RNA-seq. In contrast, BIGSI and COBS focused on indexing heterogenous *k*-mer sets, such as *k*-mers originating from various bacteria. Experiments in Bingmann *et al.* (Bingmann et al., 2019) demonstrated that SBTs like HowDeSBT could also perform well on this type of data. A trade-off exists between the construction time – in favor of COBS – and index size – in favor of the SBTs. As shown in the COBS paper, resizing BFs allows to save memory, but the latest SBT flavors also have a lightweight memory footprint because compressing BFs can achieve a similar effect as resizing. Smaller BFs also increased the false positive rate of COBS in comparison to other BF-based techniques (Bingmann et al., 2019).

It is also important to note that for many methods queries are approximate, although a number of color-aggregative methods support exact queries. Some colored DBG implementations (Vari and Vari-Merge) support additional features such as SNP and short variant discovery and graph traversal. New query types should also be considered. For instance, recording *k*-mer counts (with REINDEER) instead of presence or absence is likely to assist gene expression studies.

Color-aggregative methods and BIGSI/COBS seem better suited to query large sequences. Indeed, in these methods, the bottleneck for a single query is loading the index into memory. Then, the rest of the query consists in hashing *k*-mers, roughly in constant time per each *k*-mer. Henceforward, once the index is loaded in memory, batches of queries or large queries can be answered very rapidly. Query speeds depend on the method and its implementation. For instance, in some data structures such as the CQF in Mantis, consecutive *k*-mers are likely to appear nearby in memory, thus reducing the number of cache misses during a query. A drawback is that these structures are usually more memory consuming than SBTs. Moreover, in the case of SBTs, BIGSI, and COBS, large queries allow to mitigate the underlying BF false positive rate. SBTs and COBS do not need to load the entirety of the index into memory at query time since the query iteratively prunes irrelevant datasets. That is why SBTs are more suitable for short queries, while for large queries, *k*-mer look-ups become a bottleneck. For very large queries (e.g. the *k*-mers from a whole sequencing experiment), only AllSomeSBT (Sun et al., 2017) has an efficient specialized algorithm. The type of queries proposed by Solomon and Kingsford (Solomon and Kingsford, 2016), which has been widely adopted across SBT flavors, relies on the threshold *θ* to determine if a sequence matches a dataset. This threshold controls the sensitivity of matches with respect to sequence identity and sequencing errors. It would benefit from being further explored from a biological point of view. For instance, a single substitution in a base is covered by *k* different *k*-mers. If the indexed sequences differ from the query on that substitution, these *k*-mers will not be found in the structure, and if the value of *θ* is too high the match could be missed.

While there have been extensive empirical benchmarks to compare the performance of the different methods, analytical comparisons of their performance has been limited (see (Harris and Medvedev, 2020) for an example, though it is limited to only SBTs). The difficulty in using worst-case analysis to analyze performance in this case is that the methods are really designed to exploit the properties of real collections, and worst-case analysis is therefore not helpful. Progress can be made by coming up with appropriate models to capture the essential properties of real data and analyzing the methods using those models.

We note that there are several issues that lie outside this survey but merit mention – the selection of *k* and the presence of sequencing errors. The value of *k* is well-known to control sensitivity and specificity of the methods. Too small of a value of *k* decreases the sensitivity and too large of a value of *k* decreases the specificity. Automatically selecting the best *k* value was studied in the context of genome assembly (Chikhi and Medvedev, 2014), but to the best of our knowledge, not yet for indexing collections of read datasets. An additional related issue to this survey, is the presence or absence of sequencing errors, which result in *k*-mers that have a low number of occurrences. To address this issue, some methods (such as Cortex) filter *k*-mers with a low frequency by default, whereas others require user intervention.

The data structures surveyed in this paper should be seen as initial attempts from the community toward being able to routinely query the hundreds of thousands of samples deposited in public repositories (e.g., SRA) or private ones. An essential next step would be to have user-friendly tools. User friendliness can be seen from different perspectives. First, one may try to cast more concrete biological questions into simplified *k*-mer queries that can then be asked to the indices. Second, the results of queries could be presented in a manner that is more suitable to biologists rather than their current form, consisting mainly of the output of *k*-mer queries. For instance, a list of reads contained in the indexed datasets could be output for further investigation. However, indexing reads is more challenging, and this direction would require new developments for the structures to scale. Third, special attention given to user interfaces could help broaden the usage of these methods. Web interfaces are challenging to maintain in the long run (the group maintaining BIGSI proposed one: http://www.bigsi.io/); thus another solution could be to provide offline pre-computed indices. This way, users would only download some chunks of interest from the index for further investigation.

## ACKNOWLEDGEMENTS

This work was supported by ANR Transipedia (ANR-18-CE45-0020) and INCEPTION (PIA/ANR-16-CONV-0005). This material is based upon work supported by the National Science Foundation under Grants No. 1453527 and 1439057 to PM. This work was supported by the National Science Foundation under Grant No. 1618814. and NIH NIAID R01AI141810-01 to CB. The authors are grateful to Jan Holub for feedback on the BOSS section, and to Daniel Gautheret for his comments and corrections.

## SUPPLEMENTAL FILES

### Supplemental files outline

The Supplemental files are divided in two parts, namely, details on the methods and supplementary benchmarks. Some explanations will be given in the text below, and we provide concrete examples in Supplementary Boxes that can be found in the end of the document. First, following the main text’s organization, details on relevant *k*-mer structures (hash-based, BWT-based) and compression are given (Supplemental Boxes S1-4). Then, we give more insights about some of the set of *k*-mer sets approaches. In particular, we provide a lower-level description of structures/features that did not appear in the main document. E.g., examples of the BFT and RAMBO structures are given, as well as comparisons between specific approaches (Supplemental Boxes S5-8). Complexities are outlined in Supplemental Table S3. In the Supplemental Benchmark section, we provide the full benchmarks (Supplemental Tables S1 and S2) extracted from the different papers that led to Table 3 in the main document.

### Details on the methods

#### *k*-mer index data structures

#### Bloom filters, CQF and Othello hashing

Examples are presented in Supplemental Box S1.

#### De Bruijn graph and compacted de Bruijn graph

Instances are shown in Supplemental Box S2.

#### BOSS: BWT-based De Bruijn Graphs

The Burrows Wheeler Transform is a text transformation algorithm. It receives a sequence as input, and rearranges its characters in a way that enhances further compression. The transformation is reversible, thus the original sequence can be decoded. BOSS rearranges *k*-mers to represent the De Bruijn graph in a similar way.

Here, we briefly show how the BOSS scheme works. To begin we describe the following simple — but not space-efficient — representation of a DBG: take each unique (*k* + 1)-mer, consisting of a vertex concatenated to the label of an outgoing edge, and sort those (*k* + 1)-mers according to their first *k* symbols taken in reverse order. The resulting sorted list contains all nodes and their adjacent edges sorted such that all outgoing and incoming nodes of a given node can be identified. Thus, it is a working representation, in that all graph operations can be performed, but is far from space efficient since (*k* + 1) symbols need to be stored for each edge. Next, we show that we can essentially ignore the first *k* symbols, which will lead to a substantial reduction in the total size of the data structure.

First, we make a small alteration to this simple representation by padding the graph to ensure every vertex has an incoming path made of at least *k* vertices, as well as an outgoing edge. This maintains the fact that a vertex is defined by its previous *k* edges. For example, say *k*-mer `CCATA` has no incoming edge; then we add a vertex `$CCAT` and an edge between `$CCAT` and `CCATA`, then between `$$CCA` and `$CCAT`, and so forth. We let *W* be the last column of the sorted list of (*k* + 1)-mers. Next, we flag some of the edges in the representation with a minus symbol to disambiguate edges incoming into the same vertex – which we accomplish by adding a minus symbol to the corresponding symbols in *W*. Hence, *W* is a vector of symbols from {<u>A, C, G, T, $, -A, -C, -G, -T, -$</u>}. Next, we add a bit vector *L* which represents whether an edge is the last edge, in *W*, exiting a given vertex. This means that each node will have a sequence of zero-or-more 0-bits followed by a single 1-bit, e.g., if there is only a single edge outgoing from a node then there is a single 1-bit for that edge. Overall the representation consists of a vector of symbols (*W*), a bit vector (*L*) implemented using a rank/select Raman et al. (2002) data structure, and finally an array that records the counts of each character. It may seem surprising but these three vectors provide enough information for representing the DBG and supporting traversal operations. We refer the reader to the original paper for a detailed discussion. Lastly, we note that this representation, which is referred to as BOSS, is due to Bowe et al. Bowe et al. (2012) and was extended for storing colors Muggli et al. (2017) (see Supplemental Box S3 for an example).

##### Details on compression

To efficiently represent a *n* × *c* color matrix, over *n k*-mers across *c* datasets, different schemes have been proposed. A color class is a set of colors common to one or multiple *k*-mers. It can also be seen as a bit vector, or alternatively, a row of the color matrix. Supplemental Box S4 presents examples of the different techniques: the delta-based encoding used in Mantis+MST (a), the RRR/Elias-Fano coding (b) used e.g. in Mantis and VARI, the lossy compression using BF from Metannot (c), the BRWT principle (d), and the three strategies used in SeqOthello (e).

##### Set of k-mer sets details

###### Color aggregative methods

We first show how the different color aggregative methods combine *k*-mer sets, indexing techniques and color strategies in Supplemental Box S6.

**BFT** significantly differs from other methods: an example is shown in Supplemental Box S5. In a BFT, *k*-mers are divided into a prefix and a suffix part that are recorded in a burst trie. Prefixes are further divided into chunks, which are to be inserted into the root or inner nodes of the tree. Suffixes are in the leaves. Queries start at the tree root and progress through the path that spell the query string. In practice, each leaf stores a set of tuples: some *k*-mer suffixes along with their corresponding color classes. Bloom filters are also used in the inner nodes, to increase query speed by quickly checking the presence of a chunk.

**Supplemental Table S1.** Space and time requirements to build human RNA-seq indices. The best result for each column is shown in green. ^*a*^ refers to the HowDeSBT article, ^*b*^ to the SeqOthello article, ^*c*^ to the BIGSI article, and ^*d*^ was obtained through personal correspondence with R. Patro.

| Tool | Data Processing Time (days) | Max Ext. Memory (GB) | Time (h, wallclock) | Peak RAM (GB) | Index Size (GB) |
| --- | --- | --- | --- | --- | --- |
| SBT | 3.5<sup>b</sup> | 300<sup>a</sup> | 55<sup>b</sup> | 25<sup>b</sup> | 200<sup>a</sup> |
| AllSomeSBT | 3.5<sup>a</sup> | 600<sup>a</sup> | 25<sup>a</sup> | 35<sup>b</sup> | 140<sup>a</sup> |
| SSBT | 3.5<sup>a</sup> | 600<sup>a</sup> | 55<sup>a</sup> | 5<sup>b</sup> | 20<sup>a</sup> |
| HowDeSBT | 2.5<sup>a</sup> | 30<sup>a</sup> | 10<sup>a</sup> | N/A | 15<sup>a</sup> |
| Mantis | 130<sup>a</sup> | 110<sup>d</sup> | 20<sup>a</sup> | N/A | 30<sup>a</sup> |
| SeqOthello | 3.5<sup>b</sup> | 190<sup>b</sup> | 2<sup>b</sup> | 15<sup>b</sup> | 20<sup>b</sup> |
| BIGSI | N/A | N/A | N/A | N/A | 145<sup>c</sup> |

Note that the above description of BFT does not capture the full complexity of the data structure, and should only be used to build an initial intuition.

#### *k*-mer aggregative methods

In Supplemental Box S7, we present indexing and query details, in a similar fashion than Box 3 in the main text, but more in depth. We show the index construction and query steps in SBT, BIGSI, and show how COBS improves on BIGSI’s representation while keeping the core idea.

The false positive rate of a BF is monotonically increasing in *m/n,* where *m* is the number of *k*-mers in the dataset and *n* is the number of bits in the BF. BIGSI uses the same size *n* for all the BFs, thus the false positive rates of the BFs differ depending on how many *k*-mers are in the corresponding dataset. COBS avoid this by storing BFs of size adapted to the corresponding dataset.

Then we illustrate contrasts between the *k*-mer-aggregative methods in Supplemental Box S8. For the different flavors of SBTs, different strategies are used to store information in each node. Supplemental Box S8 shows the improvements in bitvector representation first brought by SSBT/AllSomeSBT, then by HowDeSBT. In a second Figure, BIGSI, Dream Yara and RAMBO strategies for indexing Bloom filters are compared. In the following, we outline the very recent RAMBO’s method. An example of RAMBO structure is shown in Supplemental Box S8, bottom right panel. RAMBO builds a matrix of C columns and T rows. Cells of the matrix are BFs. At construction, a given dataset is assigned to one cell per column. The corresponding BFs in those cells are each updated so that all the *k*-mers of the dataset are inserted into each of those BFs. This creates some (necessary) redundancy in the structure. Since several datasets can be assigned to a same cell, BFs become union BFs by informing for the presence/absence of *k*-mers in more than one dataset. A query is performed on the rows, each union BF giving a row-wise union of sets where the query could be present. The final sets containing the query are deduced by performing an intersection of the different set unions.

## Supplementary benchmarks

In Table S1, we show the results of different methods on a collection of 2585 of human blood, brain and breast RNA-seq datasets. This collection was first used in the original SBT paper and became a *de facto* benchmark for the other methods. It contains approximately 4 billion distinct 20-mers. Each reviewed article had its own, different, set of methods for performing a benchmark using this dataset. Here we assembled the results of three different benchmarks performed in 2018 and 2019, for which the important parameters (*k* value, abundance threshold) were identical. Even if hardware settings where different in the three studies, the presented trends (in particular, impacts on disk and RAM) remain accurate. The data processing time column refers to the time necessary to convert the original sequence files to the *k*-mer set indices (computation of Bloom filters, CQF with Squeakr, Othello). The maximum external memory column corresponds to the peak disk usage when building the index. The time column is the time required to build the set of *k*-mer sets index. The index size column is the final index size. BIGSI is not a compressed index, but the authors had explored the possibility to compress using snappy (https://google.github.io/snappy/). Parameters used for the different methods were *θ* = 0.9 andBF size of 2.10^9^ for the SBT methods, *k* = 20 as the *k*-mer size for all methods, and 34 “log slots” for Mantis from the estimation of their paper.

In Table S2 we present the space and time required to build those data structures on bacterial datasets. The table is divided into two sub-tables that correspond to two benchmarks from the literature. Contrary to the human RNA-seq experiments, these two benchmarks were not reconcilable, in terms of used datasets and parameters, thus we chose to present them separately. The first one shows results from Bingmann et al. (2019), containing 1,000 bacterial, viral and parasitic whole-genome DNA files, obtained from the BIGSI paper (http://ftp.ebi.ac.uk/pub/software/bigsi/nat_biotech_2018/ctx/). The second one is from Muggli et al. (2019) and contains 4,000 datasets 16,000 Salmonella strains (NCBI BioProject PR-JNA183844).

**Supplemental Table S2.** Space and time required to build indices on bacterial datasets. **<u>Table (a).</u>** The table shows results from a benchmark done in the COBS article Bingmann et al. (2019). The COBS benchmark contains 1,000 microbial DNA files, consisting of various bacterial, viral and parasitic WGS datasets (in the ENA as of December 2016) with an average of 3.4 million distinct *k*-mers per file. No cutoff on *k*-mer abundances was used before constructing the data structures. In the COBS benchmark, *k* was set to 31. In the table, COBS denotes for the COBS *compact* index that allows more than one batches of BFs. **<u>Table (b)</u>** shows results from the Vari-Merge article Muggli et al. (2019) The Vari-Merge benchmark contains 4,000 datasets totalling 1.1 billion distinct *k*-mers from 16,000 Salmonella strains. Note that it has more genomes than the COBS benchmark, but it possibly contains a lower variability in *k*-mer content. In the Vari-Merge benchmark, methods were run with *k* = 32, with the exception of BFT that was run with *k* = 27. When applicable, other parameters (for both Table (a) and (b)) were set to their defaults.

| Tool | Max Ext. Memory (GB) | Time (h, wall clock) | Peak RAM (GB) | Index Size (GB) |
| --- | --- | --- | --- | --- |
| **Table (a)** |  |  |  |  |
| SBT | N/A | 1.9 | 11 | 20 |
| SSBT | N/A | 8.0 | 1.5 | 3.3 |
| AllsomeSBT | N/A | 2.0 | 7.1 | 21 |
| HowDeSBT | N/A | 21 | 108 | 1.9 |
| BIGSI | N/A | 1.2 | 247 | 28 |
| COBS | N/A | 0.01 | 2.6 | 3.0 |
| SeqOthello | N/A | 0.7 | 12 | 4.4 |
| Mantis | N/A | 0.4 | 88 | 16 |
| **Table (b)** |  |  |  |  |
| Vari | 1,000 | 11 | 136 | 51 |
| Vari-Merge | 1,000 | 12 | 52 | 51 |
| Rainbowfish | 1,000 | 11 | 136 | 51 |
| BFT | 900 | 52 | 120 | 99 |
| Multi-BRWT | 1,300 | 42 | 156 | 1,300 |
| Mantis+MST | 36 | 12 | 52 | 51 |

**Supplemental Table S3.** Time complexities for the construction and query of the main approaches. *N* is the total number of distinct *k*-mers, *n* is the number of datasets, *Q* the query size (number of *k*-mers). We denote by *b* the number of bits in a Bloom filter, and *h* the number of datasets that contain at least *θ%* of *Q k*-mers. We consider *k* as a relatively small constant (around 21-63). * This time complexity is derived from Theorem 1 in Harris and Medvedev (2020) with the assumption that the size of the Bloom filter *b* is roughly 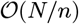. Note that there may be an additional complexity cost for building the topology of the tree through clustering. We note that in the worst case (majority of *k*-mers present in all datasets), the query complexities of SBTs would be 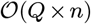.

|  | construction | query |
| --- | --- | --- |
| SBT | $\mathcal{O}(n \times b)^*$ | $\mathcal{O}(Q \times h)$ |
| VARI | $\mathcal{O}(N \times \log(N))$ | $\mathcal{O}(Q \times n)$ |
| Mantis | $\mathcal{O}(N \times n)$ | $\mathcal{O}(Q \times n)$ |
| SeqOthello | $\mathcal{O}(N \times n)$ | $\mathcal{O}(Q \times n)$ |
| BFT | $\mathcal{O}(N \times n)$ | $\mathcal{O}(Q \times n)$ |
| BIGSI | $\mathcal{O}(n \times b)$ | $\mathcal{O}(Q \times n)$ |
| RAMBO | unreported | $\mathcal{O}(Q \times \sqrt{n} \times \log(n))$ |

### Supplemental Box S1. Hashing techniques

#### Bloom filters

The example filter has a set of two functions *f* and *g.* In (1) *a* is inserted by putting 1s at positions 2 and 4 indicated by both functions. (2) *b* is inserted similarly. (3) *x* is queried, *g(x)* giving a 0 we are certain that it is not present in the filter.

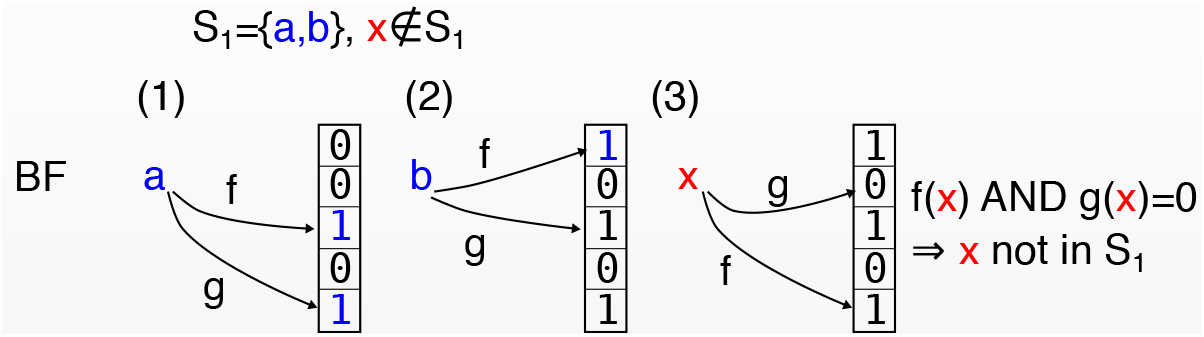

#### Counting Quotient filter intuition

Element *a* and *b* are decomposed into *a*_1_*a*_2_ and *b*_1_*b*_2_. *a*_1_, *b*_1_ are quotients, and *a*_2_, *b*_2_ are remainders. (1) During *a*’s insertion, the quotient is used to find the position of the element in the filter, and *a*_2_ is stored. The count is associated (second column). (2) similar operation for *b*. (3) *a* is re-inserted, leading to a count of 2.

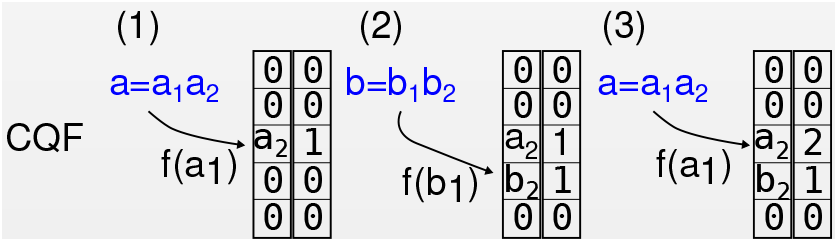

#### Othello Hashing intuition

In the example below (figure), we will focus on the case where two sets *S*_1_ and *S_2_* are hashed. A larger number of sets can be dealt with. Othello hashing uses two hash functions, denoted here by *f* and *h*. The method maintains two arrays *B*_1_ and *B*_2_, see panel (1). In the case of storing two sets, *B*_1_ and *B*_2_ are binary arrays. Elements from *S*_1_ will be mapped to one value *x*_1_ in *B*_1_ and another value *x*_2_ in *B*_2_ such that *x*_1_ = *x*_2_. Conversely, elements from *S*_2_ will correspond to identical values ((*x*_1_,*x*_2_) = (0,0) or (1,1)). In (2), the element *a* from *S*_1_ is hashed with *f* and inserted in *B*_1_ at the position given by *f* and similarly with *h* and *B*_2_. A different value will be stored in *B*_1_ and in *B*_2_ (0 and 1). The lines between those two values visually represent their association to *a.* (3) *b* is hashed the same way than *a*, ensuring again that two different values are associated to *b*. (4) Element *c* is inserted, here we cannot ensure two different values are associated to *c* without having a contradiction. Thus *b*’s 0 in *B*_2_ is modified (in red). (5) The values associated to *b* must differ, so in *B*_1_ we modify the 1 associated with *b* to a 0. (6) *x, y, z* are inserted, this time they must be associated to pairs of identical elements as they belong to *S*_2_. (7) *y* is queried by hashing it with *f* and *h* and by checking if the associated values are identical (*y* in *S*_2_) or different (*y* in *S*_1_).

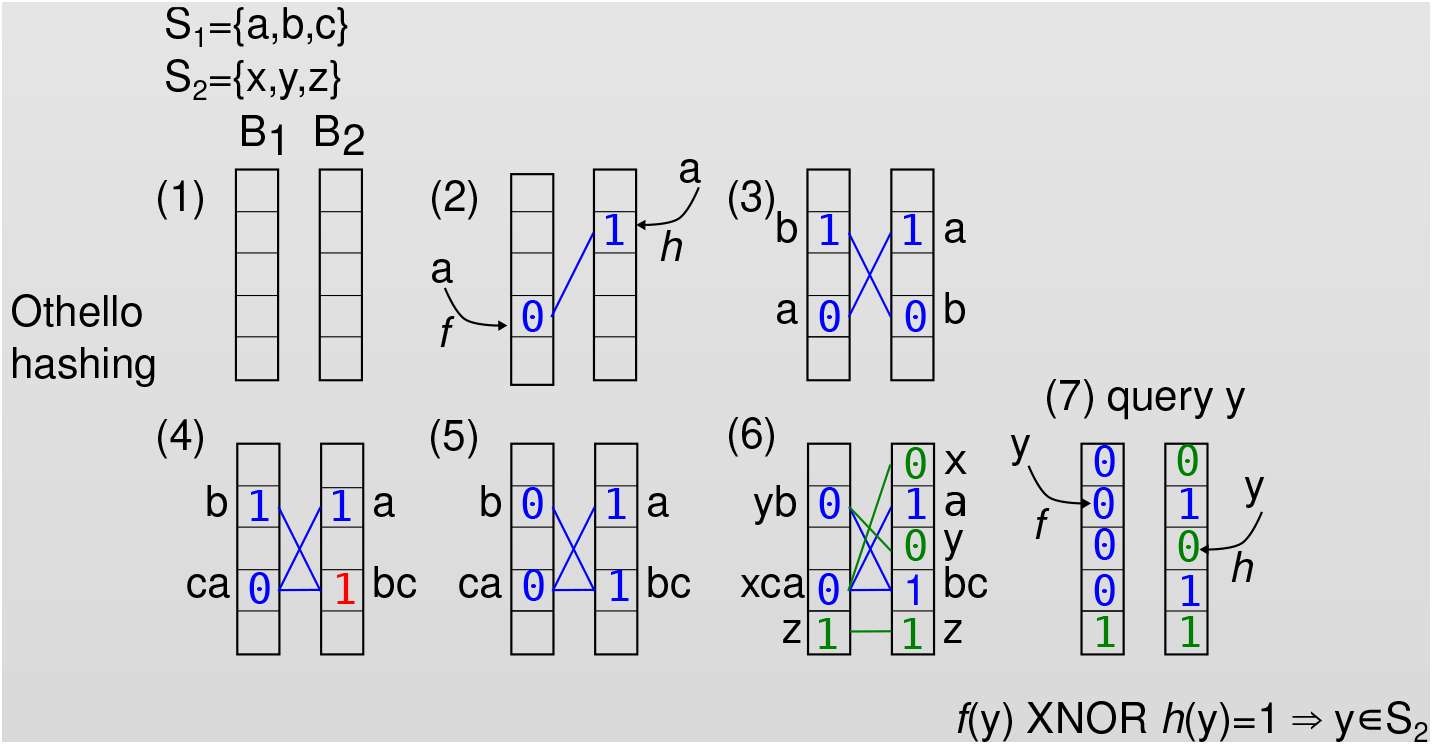

### Supplemental Box S2. De Bruijn graphs

In the example below, the first graph is a regular De Bruijn graph from the 5-mers CCTGA, ACTGA, CTGAG, TGAGA, GAGAA, AGAAC, GAACC, AACCT, AACCG. The node CTGAG has two ingoing edges and only one outgoing, GAACC has two outgoing edges and only one in-going, any other vertex in-between connecting CTGAG and GAACC has only one in-going and outgoing edge. Thus this red path can be compacted.

The second graph is the resultant <u>compacted De Bruijn graph</u>. The red path becomes a single red node, by concatenating CTGAG, **A**, **A**, **C** and **C**. It keeps the same connections than the two flanking nodes. Each resultant node is referred to as a <u>unitig</u>.

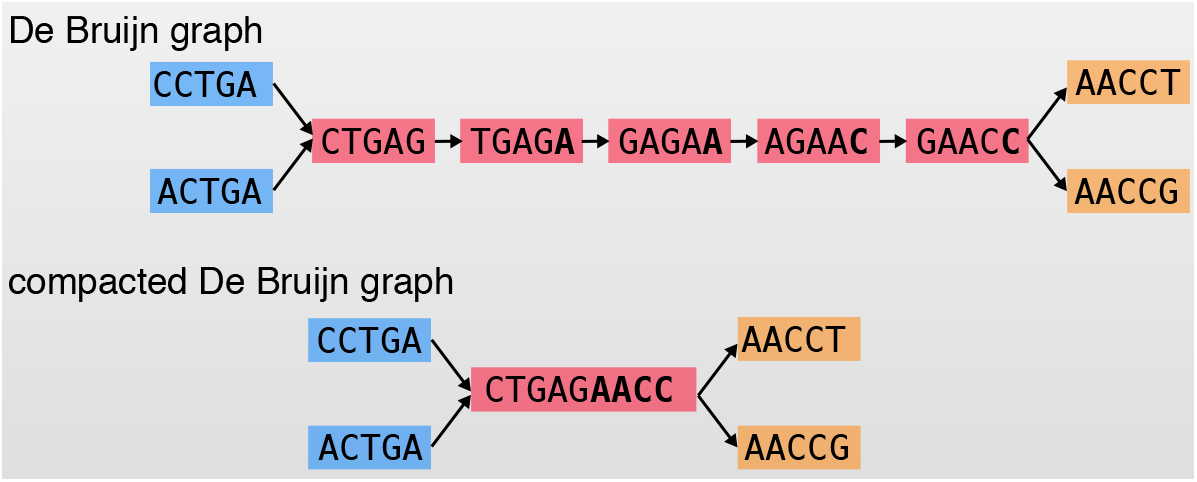

For representing the nodes, the first representation uses 5 × 9 nucleotides, and the compacted representation only uses 5 × 4 + 9 nucleotides.

### Supplemental Box S3. BOSS graph structure

We describe the BOSS data structure, as per its original flavor Bowe et al. (2012). We build a BOSS from two sequences CAGCCGA and CAGTCGA with *k* = 3. Part (2) is the De Bruijn graph from these sequences (no reverse-complements are considered here). In this representation, each vertex contains a 3-mer, and an edge represents a 4-mer existing in the original sequences, the label of the edge being the last nucleotide of this 4-mer. (3) represents the same information, but with the constraint that any nodes not containing $ must be preceded by *k* vertices (3 vertices) and must have at least an outgoing edge. Thus supplementary nodes with the padding ‘$’ symbol are added. (4) is the list of (*k* + 1)-mers in the (3) graph.

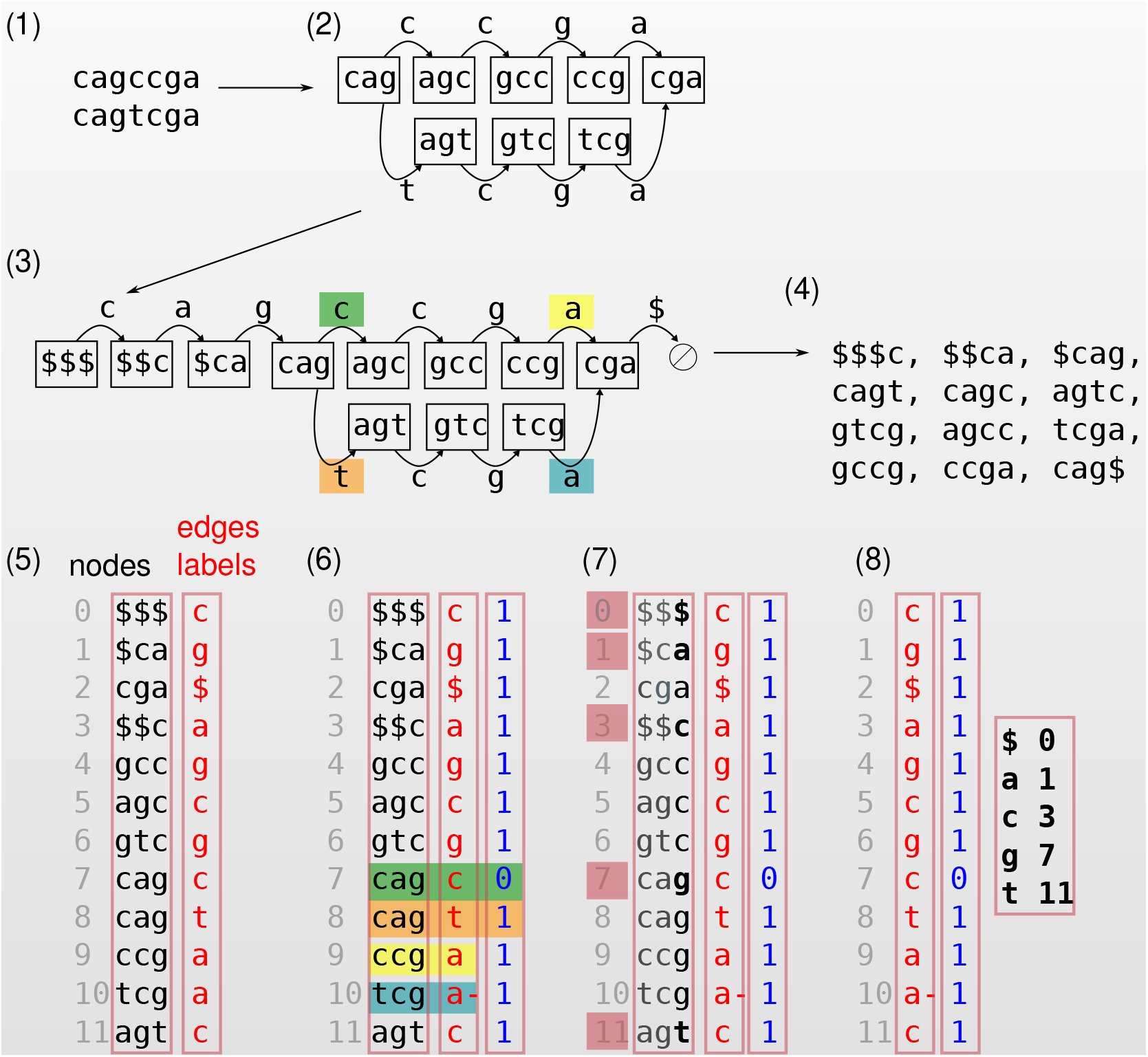

(5) These (*k* + 1) -mers are listed by lexicographic order by reading them in reverse starting from the *k*th nucleotide to the first (ties are broken by the *k* + 1-th nucleotide). This gives a matrix of nucleotides, the last nucleotide of each (*k* + 1)-mer being in a separate red column. Each line of the matrix represents a node label in the graph, and the red vector represents the edge labels. (6) In order to denote nodes that have several outgoing edges, a new vector (blue) is used. 1s indicate the last occurrence of a given node, while 0s mark its previous occurrences (they are necessarily contiguous). Here node CAG has two outgoing edges, one labeled by C (green, marked 0) and the other by T (orange, marked 1). Several edges entering the same node share the same label, all but one are marked using a –, as for yellow/blue labels. (7) Only the last column of the matrix will be kept in the BOSS. We retain the rank of each first symbol (in red): $ appears at rank 0, A at rank 1, C at rank 3,.... (8) The final information in the BOSS structure. From these tables, DBG operations such as going forward, backward from a given node are shown to be possible in Bowe et al. (2012), but we do not describe them here.

### Supplemental Box S4. Details on compression of bit matrices

Color classes can be further compressed. We present here some of the known techniques.

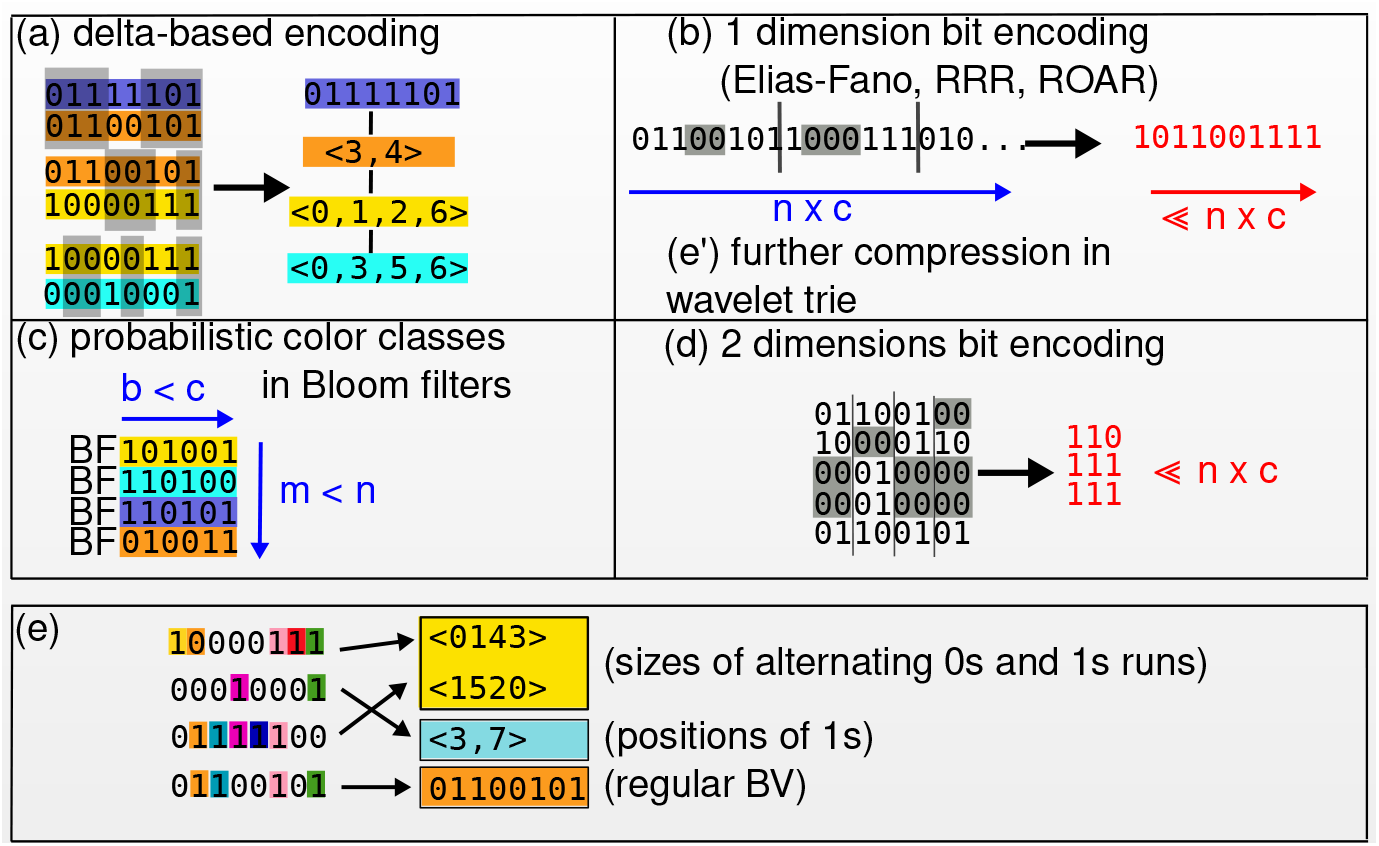

#### Delta-based encoding

(a): Differences between rows in the matrix are encoded. One can e.g. encode the (column-wise) differences between the current row and the first row as 1’s, and similarities with 0’s. This results in a sparser matrix that can be further compressed, e.g. using (b). In Figure (a), grey zones mark similarities between pairs of vectors. The purple vector is chosen as a reference, and positions of differences with the orange, yellow and blue vectors are recorded into separate lists. <u>Mantis + MST</u> uses this technique, and it also one of <u>SeqOthello’s</u> strategies.

#### Bit encoding techniques

(b,d): When encoding a color matrix: rows of the color matrix are first concatenated. The resulting bit vector is then compressed losslessly into a shorter bit vector. In (b), rows of a color matrix (not displayed) are concatenated then compressed by, e.g., finding runs of 0s (denoted in grey) and yields a smaller vector (red vector in the figure). Many tools implement this idea (<u>Vari, Mantis, Rainbowfish, BiFrost</u>, the <u>SBTs</u>). In (d), rectangular blocks of 0s (denoted in grey) in the color matrix are marked and removed. This allows 2-dimensional compression (red matrix), by storing the positions of removed blocks. This solution was proposed by <u>Multi-BWRT</u>.

Using (e), such representations can be further compressed.

#### Probabilistic color classes

(c): In <u>Metannot</u>, instead of recording the exact presence/absence of a *k*-mer within each dataset, colors are stored in a Bloom filter of size *b* < *c*. Then, the retrieval of color(s) associated with a *k*-mer becomes probabilistic (i.e., a color may be wrongly given to a *k*-mer).

#### Approximate color classes with hybrid compression

(e) Nearly identical rows are grouped, and a representative is chosen for each group. Then, depending on whether bit-vectors associated to the color classes are sparse (small amount of 1s) or dense (high amount of 1s), different compression schemes are used. SeqOthello and BiFrost use this strategy (the example shows SeqOthello’s solution). SeqOthello proposes to group similar color profiles, then uses a suitable compression technique depending on the bit-sparsity of each group. A list of integers represents the bit-vector when it has a few 1s (integers are the positions of the 1s). With many 1s, run-length encoding alternatively encodes the consecutive number of 0s and 1s. If the bit-vector has roughly the same amount of 0s and 1s, no compression is used. <u>BiFrost</u> differs a bit, by adapting different bit-encoding techniques as (b) to the different vector sparsities.

### Supplemental Box S5. Bloom filter trie

A Bloom filter trie is a tree that stores *k*-mers in its leaves. A leaf can store at most *t k*-mer, otherwise it is “burst” (i.e. transformed) into a sub-tree. The new sub-tree consists of a node *v* and two or more children of *v.* All prefixes of length *p* from the sequences in the original leaf are stored in *v*. All suffixes of length *k* – *p* that follow the *i*-th prefix in *v* are stored in the *i*-th child of *v*.

We now show how to store the following *k*-mers in a BFT: AGGCTAGCTAA, AGGCAAACTAT, AGGCTAGGATG, CTTATCCGACT, AGGTTCAGAAT, AGGCTACCCCC, with *t* = 4 and *p* = 3. In Panel (1), the first four *k*-mers can be inserted in a single leaf, since *t* = 4. (2) The fifth *k*-mer AGGTTCAGAAT (red) cannot be inserted in the leaf, requiring a burst operation. (3) To perform the burst, the prefixes of size *p* of the five *k*-mers are stored in the root. Each prefix has a pointer to its corresponding subtree. Suffixes of length *k* – *p* are stored in the leaves. (4) Inserting AGGCTACCCCC (green), it is put in the left leaf as its prefix is AGG. This induces a burst of the left leaf which is performed on Panel (5). (6) The tree is not represented explicitly, but instead, binary vectors are introduced to optimize for space. In addition Bloom filters are added in intermediate nodes to speed up queries. Note: *k*-mers are stored as tuples with their color class.

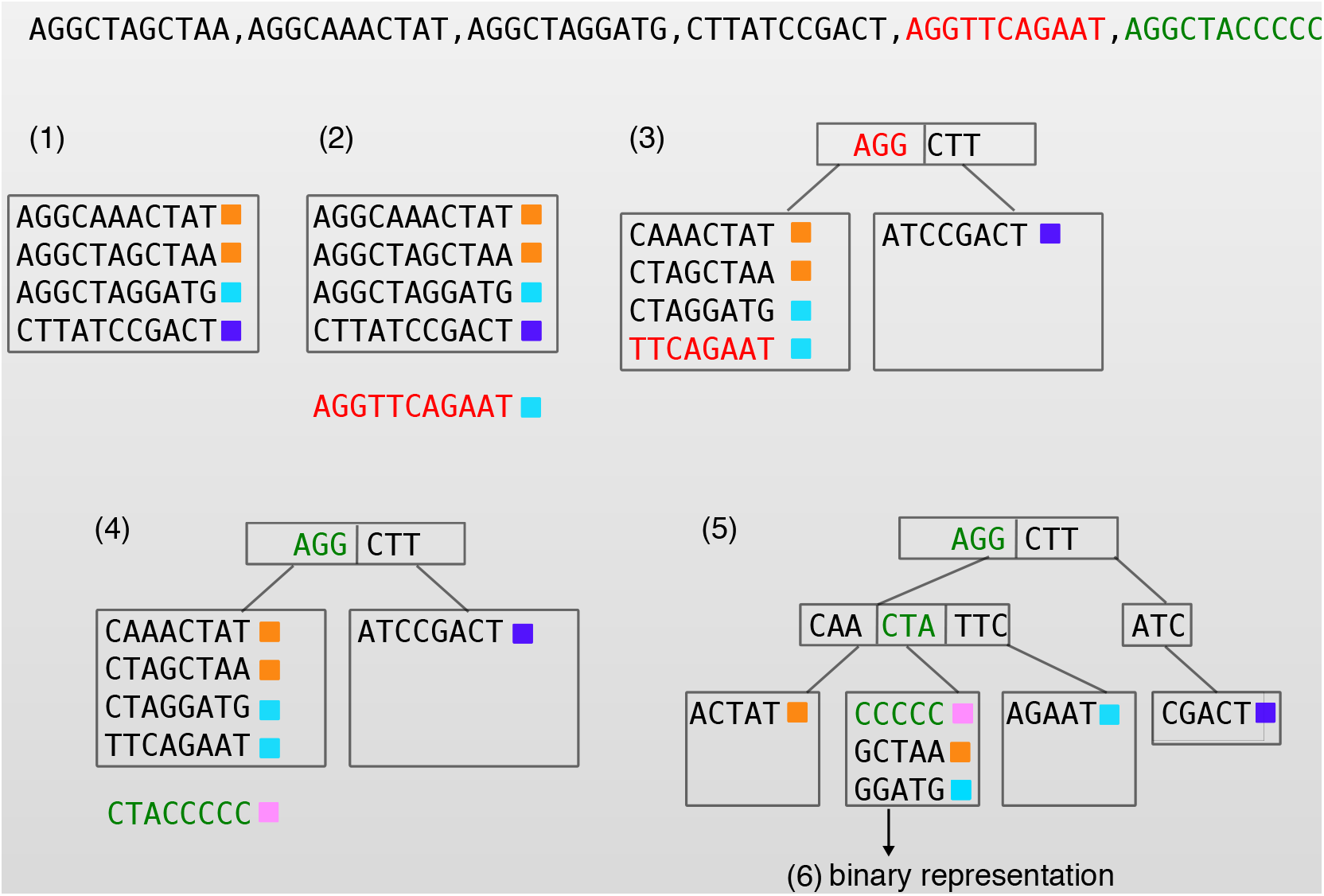

### Supplemental Box S6. Details on building blocks in color aggregative methods.

Here we give details on the color aggregative methods strategies, and in particular the *k*-mer set implementations.

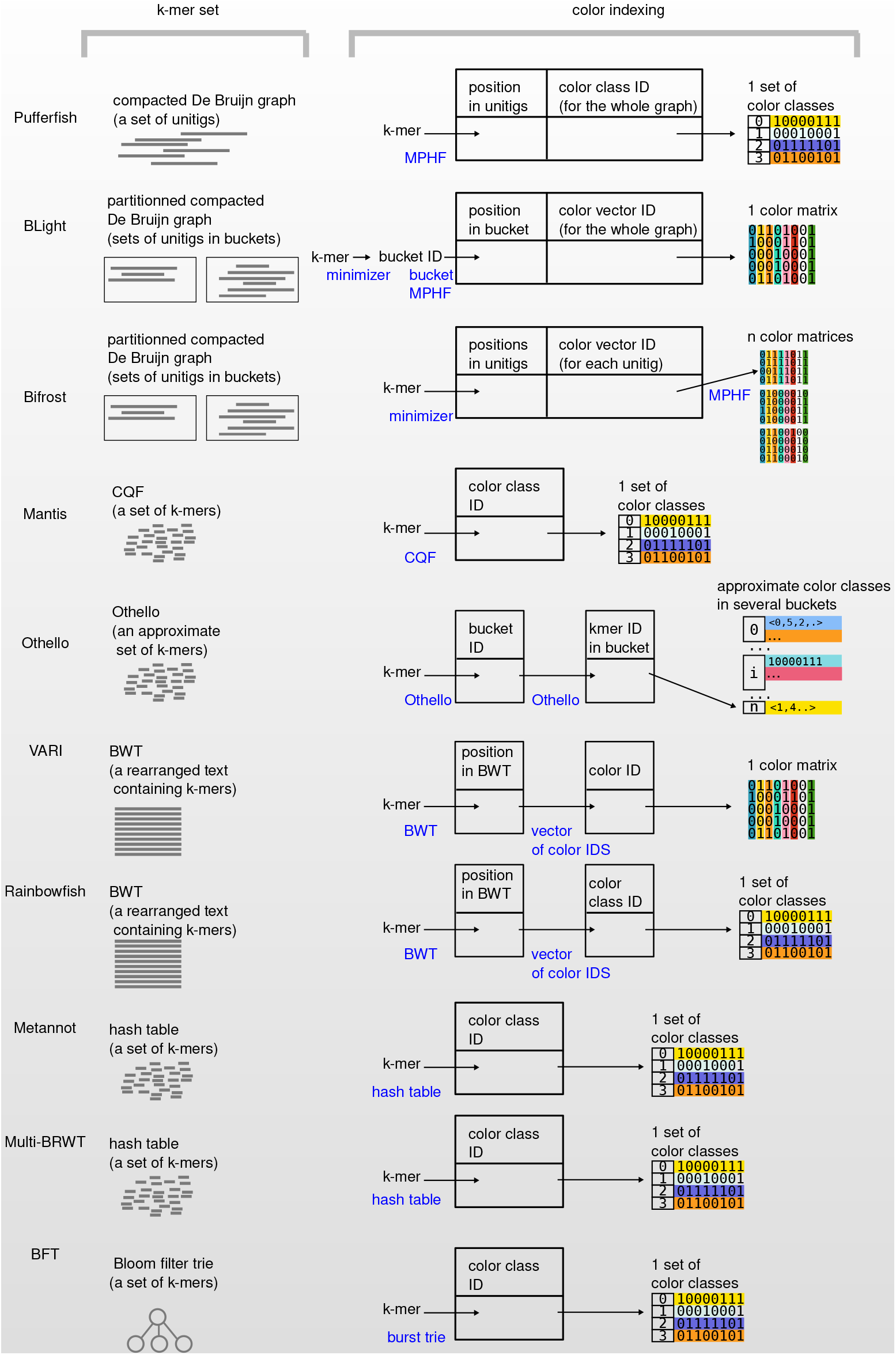

In the Figure above, the terms **compacted DBG, unitigs, BWT and MPHF** are defined in the main text.

Some of the techniques use **minimizers**. While there exist multiple definitions in the literature, here a minimizer is the smallest *l*-mer that appears within a *k*-mer, with *l* < *k*. “Smallest” should be understood in terms of lexicographical order. For example in the *k*-mer GAACT, the minimizer of size 3 is AAC, as all other *l*-mers (GAA, ACT) are higher in the lexicographical order. Minimizers are used here to create partitions of *k*-mers.Such partitioning techniques reduce the footprint of position encoding.

### Supplemental Box S7. *k*-mer aggregative methods

#### Index construction

In SBT (Panel (a) below), all BFs in leaves are have the same length, which is related to the estimated total number of distinct *k*-mers to index. The union BFs of intermediate nodes in the tree are constructed by applying a logical OR on BFs of the children BFs.

In BIGSI (Panel (b)), all BFs must also have the same size. COBS uses the same principle but Bloom filters do not all have the same size.

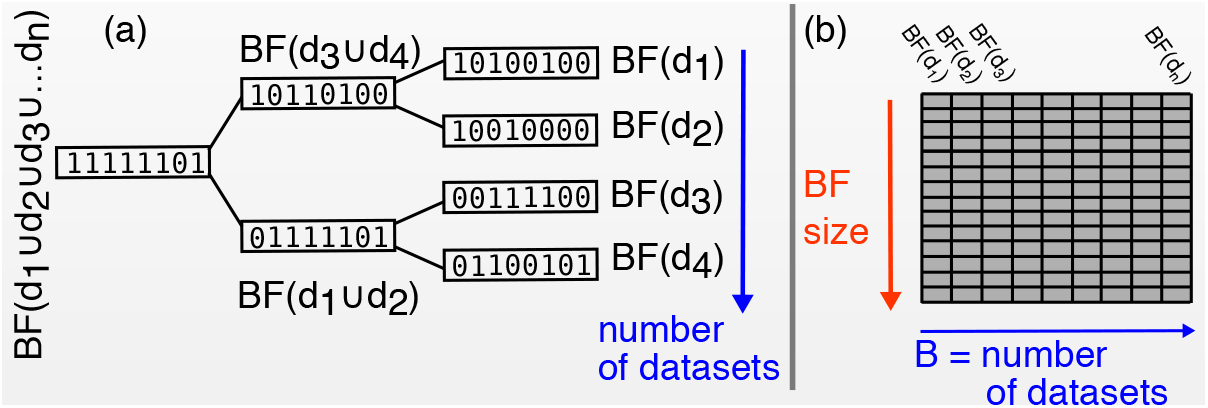

#### Queries

Here we show in more details how queries are performed in SBTs and BIGSI (see Box 3 in the main text for a more abstract sketch). In the Figure below (left figure), we consider a single hash function *f*, as it was presented in HowDeSBT’s paper for instance, and *θ = 2/3.* For BIGSI (right figure), we present the query step for one *k*-mer and three hash functions (*f*, *g, h).* Note that usual queries are composed of more than one *k*-mer and aggregate the *k*-mers results. A given *k*-mer is hashed, leading to one or several rows to lookup. In Figure (b) below, the query is performed on the green rows. Each queried row informs on the datasets that may contain the query *k*-mer. All the returned bit vectors are then summarized vertically into a single vector, using a logical AND operation. Positions yielding 1s after this operation correspond to datasets that contain the *k*-mer (in the Figure example, the *k*-mer is present only in dataset 1). (c) The same principle is used when matrices of several sizes contain the BFs. Hashes are simply adapted to the different sizes using a modulo.

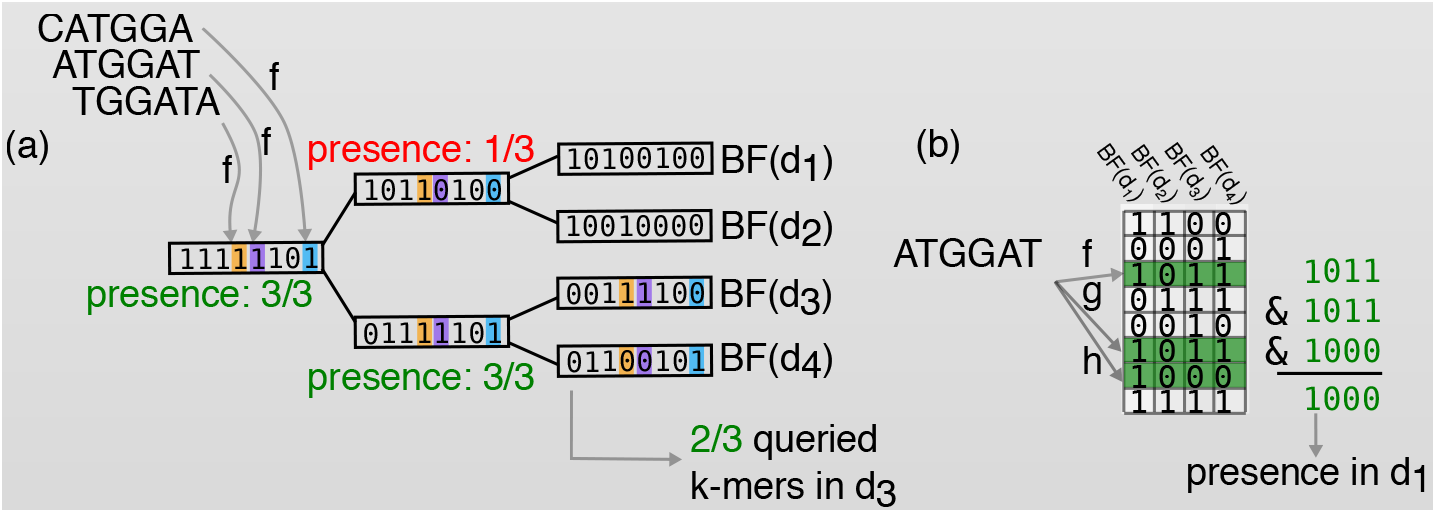

### Supplemental Box S8. Advanced and alternative k-mer aggregative methods

#### Comparison between SBT approaches

For SBTs, different strategies are used to store information in each node. In the example below, the first level of each node is a plain Bloom filter, as used in the original SBT approach. The second level is the *how + det* representation used in HowDeSBT. The third is the equivalent *all + some* or *sim + rem* representations used by AllSomeSBT and SSBT. The three approaches are shown in four nodes.

At some positions (marked in green), the bits will have the same value across all the nodes of the subtree. Those bits are marked as *det* (determined), and when they are, the *how* field records their values. In the *sim + rem* and *all + some* systems, such bits will have a value set to 1 in *all/sim* in the root node if and only if they are determined as 1. In this node, the *some/rem* vector stores values such that *all* ∪ *some = BF* or *sim* ∪ *rem = BF*, where *BF* is the Bloom filter of the node.

At the second level of the tree, new bits become determined (orange in the left subtree and blue in the right subtree). The same rules apply. Moreover, bits that were marked in the upper levels are non informative (red boxes). They can be removed from the structure, but their positions are recorded using an auxiliary bit vector (not shown).

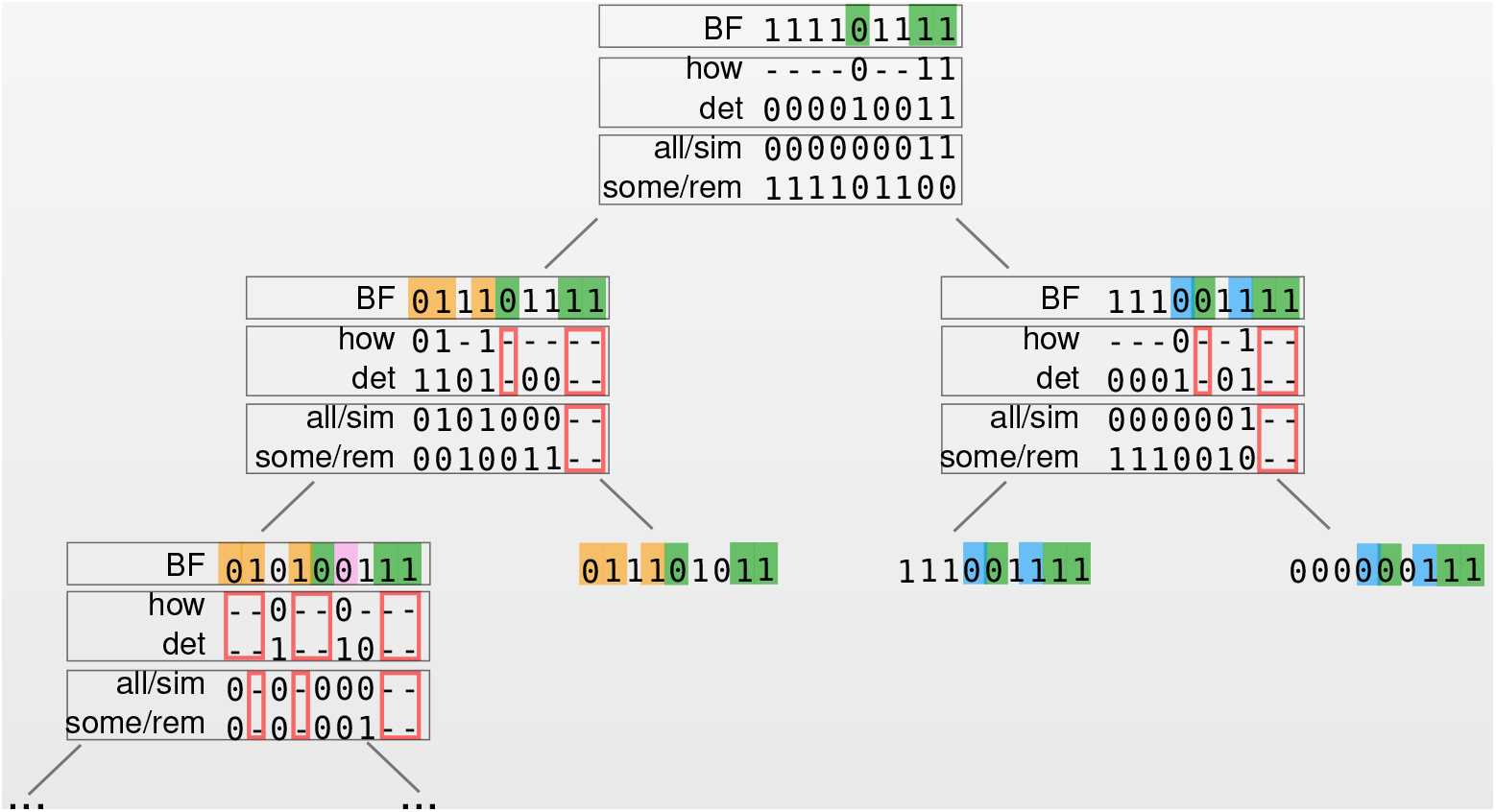

#### Comparison between Bloom filter approaches

Bloom filters are grey panels in all figures below. Left: BIGSI and COBS differences reside in the column sizes. COBS queries are as fast as BIGSI, using a system of modulos with the different BF sizes. Middle: the DREAM-Yara index is built by interleaving the bits of each Bloom filters. Bits of the same rank are grouped together in bins of size *n.* Right: each element in the RAMBO matrix is a Bloom filter (each column of the matrix stacks complete BFs). Contrary to SBTs, RAMBO merges randomly the datasets. Queries in SBTs are top-bottom, RAMBO queries each row and uses the intersection result.

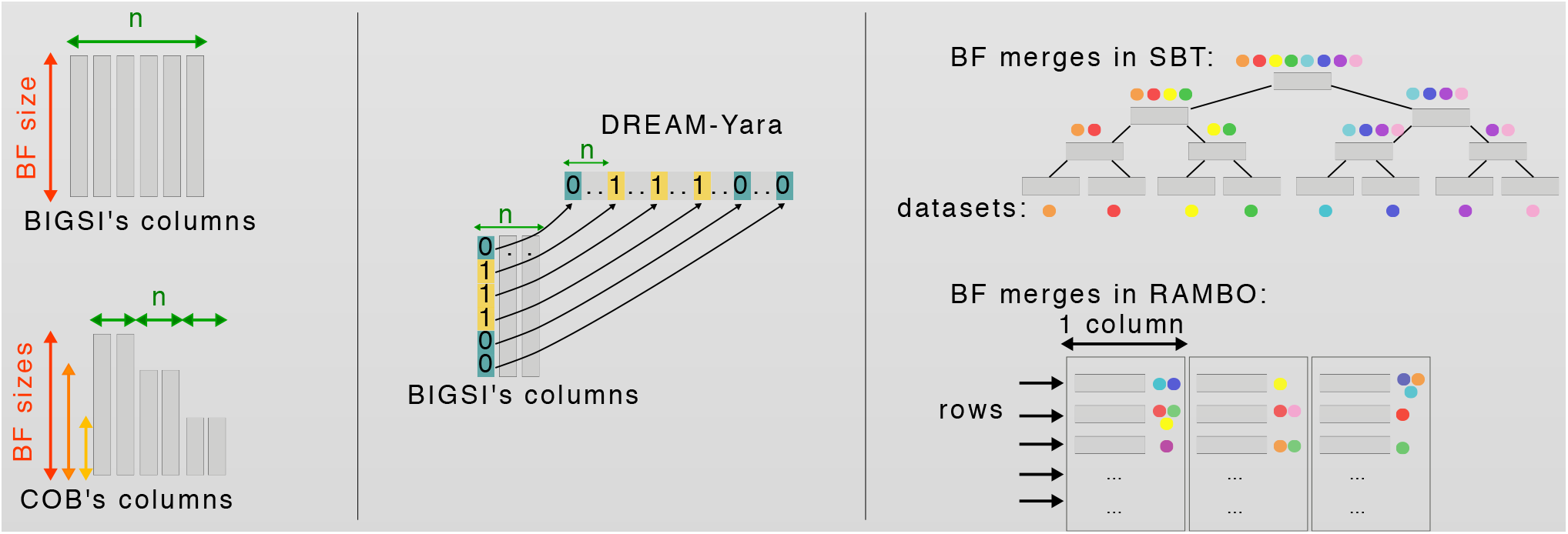

